# Targeting Epsins to Inhibit FGF Signaling while Potentiating TGF-β Signaling Constrains Endothelial-to-Mesenchymal-Transition in Atherosclerosis

**DOI:** 10.1101/2022.08.09.503324

**Authors:** Yunzhou Dong, Beibei Wang, Mulong Du, Bo Zhu, Kui Cui, Siu-Lung Chan, Douglas B. Cowan, Sudarshan Bhattacharjee, Dan Shan, Scott Wong, Joyce Bischoff, MacRae F. Linton, Hong Chen

## Abstract

**BACKGROUND:** Epsin endocytic adaptor proteins are implicated in the progression of atherosclerosis; however, the underlying molecular mechanisms have not yet been fully defined. In this study, we determined how epsins enhance endothelial-to-mesenchymal transition (EndoMT) in atherosclerosis and assessed the efficacy of a therapeutic peptide in a preclinical model of this disease.

**METHODS:** Using single cell RNA sequencing (scRNA-seq), combined with molecular, cellular, and biochemical analyses, we investigated the role of epsins in stimulating EndoMT using knock-out mouse models. The therapeutic efficacy of a synthetic peptide targeting atherosclerotic plaques was then assessed in *Apoe*^−/−^ mice.

**RESULTS:** ScRNA-seq and lineage tracing revealed that epsins 1 and 2 promote EndoMT, and the loss of endothelial epsins inhibits EndoMT marker expression as well as transforming growth factor-beta signaling *in vitro* and in atherosclerotic mice, which is associated with smaller lesions in *Apoe^−/−^* mouse model. Mechanistically, the loss of endothelial cell epsins results in increased fibroblast growth factor receptor-1 (FGFR1) expression that inhibits TGF-β signaling and EndoMT. Epsins directly bind ubiquitinated FGFR1 through their ubiquitin-interacting motif (UIM), which results in endocytosis and degradation of this receptor complex. Consequently, administration of a synthetic UIM–containing peptide API significantly attenuates EndoMT and progression of atherosclerosis.

**CONCLUSIONS:** We conclude that epsins potentiate EndoMT during atherogenesis by increasing TGF-β signaling through FGFR1 internalization and degradation. Inhibition of EndoMT by reducing epsin-FGFR1 interaction with a therapeutic peptide may represent a novel treatment strategy for atherosclerosis.

## INTRODUCTION

Atherosclerosis is one of the most challenging vascular diseases to cure, and it has been causally linked with heart attacks and strokes. Generally, the disease involves multiple cell types, including endothelial cells, vascular smooth muscle cells, immune cells, and others^1^. One of the major pathological features of atherosclerosis is the accumulation of arterial plaques that can progress from simple fatty streaks to complex lesions that can reduce blood flow or rupture leading to thrombosis and ischemic cardiovascular events ^1, 2^. At the tissue level, each cell type can contribute to atherosclerosis; however, the endothelium has been proposed to be crucial for the initiation and progression of atherosclerosis^3, 4^.

Chronic vascular inflammation is a hallmark of atherosclerosis^5^ and induces endothelial-to-mesenchymal transition (EndoMT)^6^. During EndoMT, vascular smooth muscle cell (VSMC) or mesenchymal stem cell-like markers, such as α-SMA, SM22α, Collagen 1a, Fibronectin, and N-cadherin, are expressed in endothelial cells^7, 8^. This process is thought to allow mesenchymal transition of non-SMC-derived cells that are capable of maintaining indices of lesion stability; however, this temporary protection is lost when atherosclerosis progresses to a more advanced stage, presumably because of increased inflammation associated with the advanced lesion^9^. Furthermore, other studies suggest that EndoMT exacerbates vascular inflammation in atherosclerosis^10, 11^.

It is known that TGF-ß signaling is a primary driver of EndoMT^12, 13^, while disturbed shear stress and cytokines are the most common activating modulators for TGF-ß signaling^11^. In contrast, FGFR1 signaling has been proposed as an endogenous inhibitor that suppresses the TGF-ß pathway^7, 11^. Using an endothelial lineage tracing approach supported by cell-specific marker staining^10, 14^, EndoMT has been linked to multiple cardiovascular^15–17^ and metabolic diseases, including atherosclerosis, obesity, hypertension, hyperlipidemia, and diabetes^7^. As a result, EndoMT has been suggested to be a target for treating vascular diseases^11, 18–20^, including atherosclerosis.

Epsins are ubiquitously-expressed adaptor proteins involved in the regulation of the endocytosis^21, 22^ of plasma membrane receptors complexes^23–27^, which facilitates their degradation to influence endothelial cell signaling^23, 25–27^. Using epsin mutant mouse models, we have discovered that these proteins regulate Notch, VEGFR2, and VEGFR3^23–28^ by binding to ubiquitinated membrane receptors to physiologically or pathologically modulate cell function in embryogenesis^28^, angiogenesis, lymph angiogenesis^23, 25–27^. Our previous reports showed epsins promote atherosclerosis in the endothelium and in macrophages.

Deletion of epsins in endothelial cells attenuates atherosclerosis by stabilizing IP3R1 degradation^29^ and by blunting inflammatory signaling through TLR2/4^30^ in the *Apoe*^−/−^ atherosclerosis mouse model, suggesting that epsins promote atherosclerosis by modulating inflammatory signaling pathways. Macrophage-deficient epsins also reduce atherosclerosis by binding to the LDL Receptor-related Protein 1 (LRP-1) to increase efferocytosis^31^. Loss of epsins in macrophages decreases pro-inflammatory M1 macrophages and increases anti-inflammatory M2 macrophages^31^. These observations suggest that epsins promote atherosclerosis by regulating inflammation.

To investigate the role of epsins in EndoMT, we generated inducible mutant mice lacking epsins 1 and 2 in endothelial cells and bred these mice into an atherosclerotic (*i.e., Apoe*^−/−^) background^29^. Here, we show that epsins are required for EndoMT, and that the loss of these proteins in the endothelium reduces EndoMT by permitting sustained FGFR1 signaling by inhibiting the degradation of this receptor complex. Importantly, we demonstrate the efficacy of blocking epsin-FGFR1 interactions specifically in atheromas using systemic administration of a targeted epsin UIM-containing peptide to inhibit EndoMT and atherosclerosis progression; thereby, providing a new approach to combat this disease.

## METHODS

ScRNA-seq datasets have been deposited in a publicly available repository as indicated below. All other data that support the findings of this study are available from the corresponding authors on reasonable request.

### Genetic mouse models

Mice used in this study are C57Bl6/J genetic background. C57Bl6/J strain is defined as wild type (WT). EC-specific deletion of epsin was established by crossing Epsin 1^fl/fl^: Epsin 2^−/−^ mice with endothelial cell (EC)-specific Cre transgenic iCDH5-Cre mice, and these mice were bred with the *Apoe*^−/−^ atherosclerotic mouse model and backcrossed seven times. Mice with an inducible deletion of endothelial Epsins 1 and 2 on an *Apoe*^−/−^ background (EC-iDKO/*Apoe*^−/−^) were fed a Western Diet (WD) for 12 to 14 weeks beginning at 8 weeks of age^32, 33^. Aortic roots and hearts were isolated and processed for evaluation of atherosclerotic lesion as previously described^29, 31–33^. Lesion area was quantified using NIH Image J.

### Evaluation of EndoMT markers by immunostaining *in vivo* and *in vitro*

*Apoe*^−/−^ mice, EC-iDKO/*Apoe*^−/−^, control, or API peptide-treated mice were evaluated by co-staining CD31 with EndoMT markers, including αSMA, MyH11, SM22, and Calponin. EndoMT is expressed by the overlapping percentile (%) of CD31 with VSMC markers. For *in vitro* EndoMT marker staining, mouse aortic endothelial cell (MAEC) cultures were treated with TGFβ (10 ng/mL) as indicated.

### Aortic sample preparation for single cell RNA sequencing (scRNA-seq)

Mouse aortas were cut into fine pieces and placed in 6.5 mL enzyme solution at 37°C for 1.5 hours (5 mL DMEM, 1 mL trypsin, collagenase type I 10 mg/mL, collagenase type IV 10 mg/mL, 50 µL elastin (2.5 mg/mL stock), collagenase type II 200 µL (stock: 1 mg/mL), Collagenase XI 200 µL (stock: 1 mg/mL), Hydrolase 200 µL (stock: 1 mg/mL), Liberase 100 µL (stock: 3.85 mg/mL, Roche), DNase I 10 µL (stock: 5U/µL, Sigma). This solution was then filtered using a 40 µm strainer (Falcon, NC), centrifuged (400xg for 5 min) and the cell pellet was suspended in 90 µL FEB buffer containing 0.5% BSA and 2 mM EDTA in PBS. 10 µL of CD31 Microbeads (MACS) was added to the suspension at 4°C for 15 min. Microbeads were passed through a LS column (MACS), Cells were processed for scRNA-seq using the 10xGenomics platform and sequencing was performed by BGI-HK.

### Data analysis of single cell RNA seq

We used the Cell Ranger and R package Seurat v.4.0.2^34^ to perform scRNA-seq analysis. Briefly, we generated a normalized gene expression matrix harboring a total of 5,016 and 5,521 cells for EC-iDKO and WT mouse models, respectively, which were then integrated using multiple canonical correlation analysis (CCA). Two-dimensional Uniform Manifold Approximation and Projection (UMAP) and Graph-based clustering were used for visualization. To validate our results, we empirically introduced well-defined markers, including Pecam1, Cdh5, Cldn5, Tek, and Notch1 for EC cells and Acta2, Tagln, Cald1, Mylk, and Col1a1 for VSMC cells, to explore cell types from cell clusters. Differential gene expression between groups of interest was performed using FindMarkers function implemented in Seurat. Genes with log2 fold change > 0.5 and P < 0.05 were considered differentially expressed. Pathway enrichment analysis for group comparison was performed via R package ReactomeGSA (https://github.com/reactome/ReactomeGSA) using the ‘analyze_sc_clusters’ function. The trajectory inference for single cell data from EndoMT models was performed via R package slingshot [PMID: 29914354]^35^.

### Lineage tracing

TdTomato mice were bred to WT or EC-iDKO mice to create WT: TdTomato and EC-iDKO: TdTomato mice, and these mice were then bred to iCDH-cre mice to establish WT: TdTomato: iCDH-cre and EC-iDKO: TdTomato: iCDH-cre mice. Tamoxifen was given to these mice seven times to delete epsins and activate the expression of the TdTomato gene. At 8 weeks of age, these mice were fed a WD for 8 weeks. Aortic roots and arches were fixed and immunostaining with CD31 prior to image acquisition using a confocal microscope (LSM880, Carl Zeiss).

### Administration of synthetic peptides (Control and API) to *Apoe*^−/−^ mouse model

(1) Prevention protocol: Eight-week-old male *Apoe*^−/−^ mice were fed a WD for eight weeks and injected with Control or API peptides. The dose of each peptide was 25 mg/kg, with two injections per week, I.V. for two months.

(2) Therapeutic protocol: Eight week old male *Apoe*^−/−^ mice were fed a WD for 6 weeks, and split into two groups: control or API peptides. Peptides were administered as described above for 6 weeks. The dose of each peptide was 25 mg/kg, with two injections per week, I.V. injection.

### Cell migration assay

A Boyden chamber assay protocol was used in this study to assess cell migration as modified from Wang et al^36^. In brief, we isolated MAECs from WT and EC-iDKO mouse aortas as previously described^29^. Cells were treated by 5 µM tamoxifen for 4 days to delete epsins, followed by the 10 ng/mL TGF-ß treatment for another 4 days. Cells were trypsinized and counted by microscopy. 750 µL test medium with or without 10% FBS was added in the 24-well plate wells, and 50K cells in 500 µL DMEM were added into the upper chamber. The plate was kept in the 37°C CO_2_ incubator for 7 hours. The non-migrated cells on the upper side of the membrane were removed using a cotton swab. Cells were then fixed by methanol and stained by eosin^36^. Images were captured using inverted microscopy (Axio Observer, Zeiss).

### Cell surface FGFR1 biotinylation and western blot analysis

FGFR1 biotinylation was performed as described previously for VEGFR2^25^. Biotinylated surface FGFR1 was detected by western blotting using an anti-FGFR1 antibody and transferrin served as internal control.

### Standard methods

Isolation of primary mouse aortic endothelial cells (MAECs) and generation of epsin-deficient cells (DKO) *in vitro* were performed as previous report^29^. Western blotting, Immunoprecipitation (IP) analysis, immunostaining, and cell culture were performed according to routine methodologies in our laboratory^23, 25^. Plasmids or siRNA transfection, electroporation, and RT-PCR were performed as described^23, 25, 27^.

### Study approval

Animal studies were approved by the Institutional Animal Care and Use Committee (IACUC) at Boston Children’s Hospital.

### Statistical analysis

Statistical analysis was conducted using Prism 8.0 (GraphPad Software). Data are presented as mean ± SEM. Data were analyzed by two-tailed, unpaired or paired Student’s *t-*test or ANOVA A *p* value of less than 0.05 was considered statistically significant.

## RESULTS

### Single cell RNA sequencing revealed epsins play a critical role in EndoMT

To investigate whether epsins play a role in EndMT, we performed scRNAseq analysis. WT and EC-iDKO mice were fed western diet (WD) for 14 weeks, followed by the isolation of aortas and enzymatic digestion (Supplemental Figure 2; Supplemental Methods) and scRNA seq. Our data show that the two groups of cells from WT and EC-iDKO were well distributed and merged using Principal Component Analysis (PCA) and Uniform Manifold Approximation and Projection for Dimension Reduction (UMAP) plot (Figure 1A & B). We then divided the cell population into 10 clusters. Again, the UMAP distribution in WT and EC-iDKO group are similar, comparable and nonbiased (Figure 1C). Together, all these data suggest our data quality control is appropriate.

**Figure 1.**
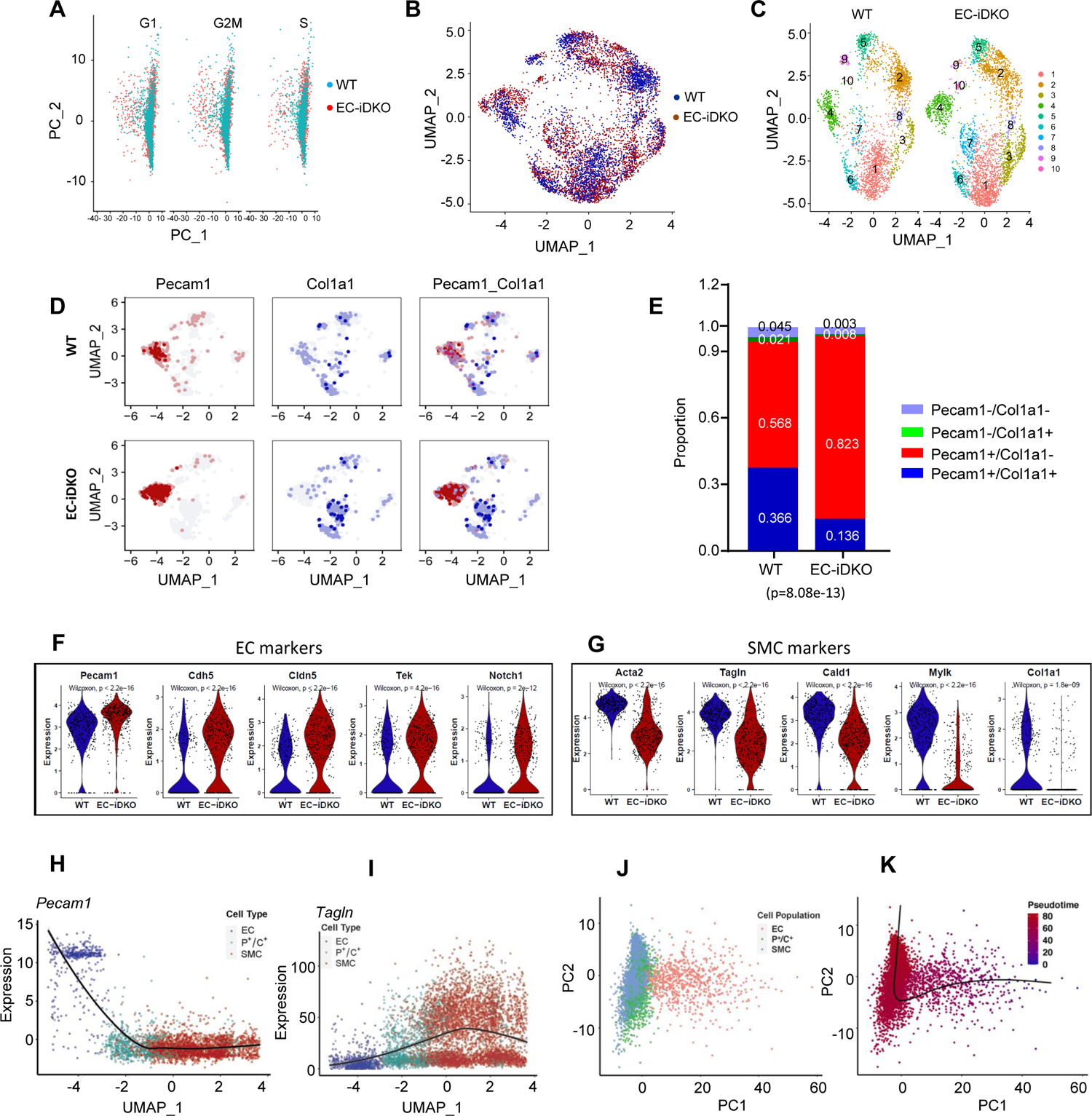
Single cell RNA sequencing analysis reveals epsins are required for EndoMT. **A**, **B**, Data quality control for WT and EC-iDKO cells in PCA and UMAP plots; **C**, WT and EC-iDKO cells were clustered to 10 groups in UMAP visualization; **D**, Representative makers for the labeling cell distribution in UMAP visualization for EndoMT models. *Col1a1* is specific to SMC and *Pecam1* to EC. SMC cells are excluded in consideration of the cell origin from the mouse blood vessels. **E**, EC enriched cells (cluster 4) were further divided to 4 subgroups as indicated. The proportion of EC and Endothelial-to-Smooth Muscle Transition Cell (EST) types was identified (*p*=8.08e-13 by χ^2^ test). **F**, **G**, Violin plot of EC and SMC markers differentially expressed in WT and EC-iDKO in the endothelial subgroups of five EC marker positive cells (*Pecam1, Cdh5, Cldn5, Tek*, and *Notch1*) and five SMC maker positive cells (*Acta2, Tagln, Cald1, Mylk,* and *Col1a1*). *P* value is calculated via Wilcoxon test. **H**, **I**, Gene expression tendencies of representative markers specific to EC (*Pecam1*) and SMC (*Tagln*) through EC-P^+^/C^+^-SMC axis of EndoMT shown as normalized expression versus UMAP_1 (also refer to the Supplemental Figure 4 with more maker genes). **J**, **K**, Inference of pseudo-times for EndoMT trajectory from EC to SMC by *Slingshot*. A linear transition was identified from EC to SMC in aortas of mouse models.

We then analyzed established markers of endothelial and VSMC, and we found that five EC markers Pecam1, Cdh5, Cldn5, Tek and Notch1 were all exclusively expressed in cluster 4 in the EC-iDKO population. The same was observed in WT mice, except Pecam1, which was also expressed in group 5-10 (possibly immune cells) (Supplemental Figure 3); this data strongly suggests that cluster 4 is the EC enriched population. The SMC specific marker genes are all expressed in 1-10 groups in EC-iDKO and WT population with different expression levels (Supplemental Figure 3).

Since clusters 1-3 do not express any EC marker genes but only express SMC marker genes, these three groups are defined as SMC or SMC-like cell population (Supplemental Figure 3). We then use Pecam1 and collagen a1 (Col1a1) to pinpoint EC and SMC marker population in UMAP (Figure 1D), and it is obvious that these two clusters have overlapping cells that co-express endothelial and SMC markers; we defined this group of cells as P^+^/C^+^ cells. The proportion of Pecam1 and Col1a1 is plotted in Figure 2E (*p*=8.08e-13). For the Pecam1 positive cluster, we further divided it into 4 groups by Pecam1 and collagen 1a1 (Col1a1, SMC marker) as displayed in Figure 1E, in which Pecam1^+^/Col1a1^−^ population or endothelial cells is 1.45 fold in EC-iDKO versus WT, a 45% increase in EC-iDKO mice; while the P^+^/C^+^ cells in WT is 2.7 fold increase compared to EC-iDKO, suggesting that more ECs are becoming P^+^/C^+^ cells in WT, while loss of epsins in endothelium significantly blocked the process of EC transforming towards SMC.

**Figure 2.**
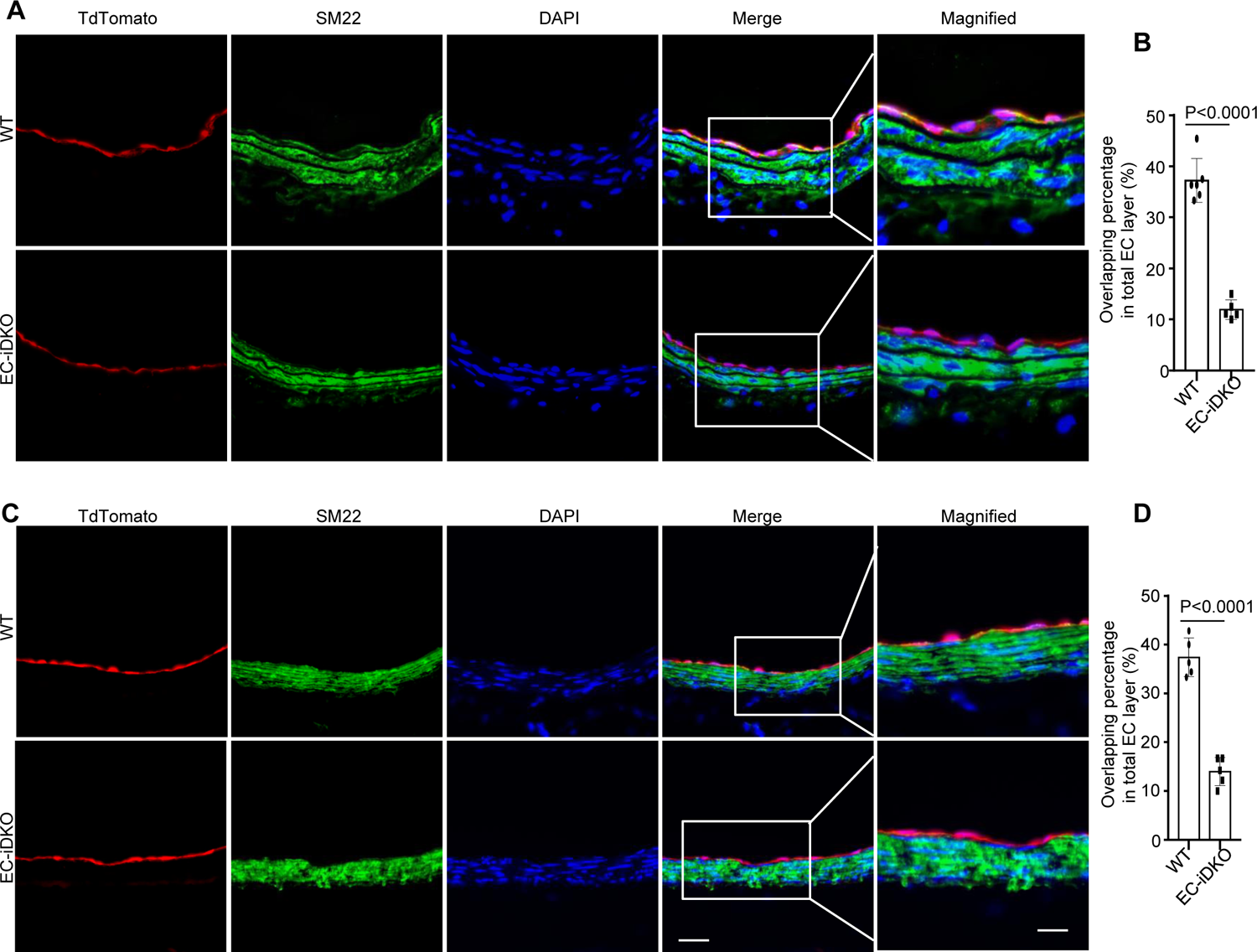
Mouse lineage tracing suggests that epsins are crucial to promote EndoMT. TdTomato mice of WT and EC-iDKO were all injected tamoxifen seven times and fed WD for 8 weeks. EndoMT marker SM22 was compared in BCA (brachiocephalic artery) (**A, B**) and aortic root (**C, D**) and quantified. n=5, *, ** P<0.001. Scale bar: 20 µm, and magnified: 10 µm.

This result is supported by measuring specific EC and SMC markers in Pecam1+ cluster. As shown in Figure 1F & 1G, violin plots show that EC markers (Pecam1, Cdh5, Cldn5, Tek and Notch1) are significantly upregulated in EC-iDKO aortas than that in WT aortas, while SMC markers are downregulated in EC-iDKO mice; importantly, Col1a1 is exclusively expressed in WT but seldom expressed in EC-iDKO mice. Together, loss of epsins from the endothelium significantly blocks the EndMT.

### Trajectory of EC to SMC differentiation in WT and EC-iDKO mice subjected to inflammatory stimuli

As displayed in Figure 1H & I, as well as Supplemental Figure 4H, the trajectory plot clearly demonstrated the transition of EC to P^+^/C^+^ and SMC process by EC marker Pecam1 and SMC marker Tagln in WT and EC-iDKO cell population. Similarly, all the tested EC and SMC markers showed the similar tendency by using integrated population of WT and EC-iDKO cells, (Supplemental Figure 4B & C).

To mimic the EndMT with the “time” change, we plotted EC, P^+^/C^+^ cells and SMC in PCA plot, and performed pseudotime analysis. As suggested by Figure 1J & K, the three populations clearly clustering, overlapping and showed a typical different cell type distribution in pseudotime axis, suggesting EC→ P^+^/C^+^ →SMC cell type transition.

### TdTomato lineage tracing shows epsins are crucial to promote EndoMT

We demonstrated that oxLDL stimuli and WD-fed *Apoe*^−/−^ mice upregulate epsin expression in endothelium in our previous report^29^. EndoMT has been positively linked in human atherosclerosis^7^. These results imply that epsins are associated with EndoMT in atherosclerosis. To further test our hypothesis, we bred the transgenic TdTomato mice to EC-iDKO and WT mice to label the endothelium in the mice of WT:TdTomato: iCdh5-Cre and EC-iDKO:TdTomato:iCdh5-Cre. All mice were given seven tamoxifen injections to activate the Tdtomato gene expression for the labeling of endothelial cells, followed by WD feeding for 8 weeks. EndoMT marker SM22α was immunofluorescently stained in BCA (brachiocephalic artery) and aortic roots of WT and EC-iDKO mice. As shown in Figure 2, in the BCA of arch (A, B) and aortic roots (C, D), TdTomato gene expression (red color) co-localized with SM22α in EC-iDKO mice is significantly reduced when compared to WT mice, suggesting that endothelial epsins are necessary to promote EndoMT.

### Loss of endothelial epsins attenuates EndoMT in atherosclerotic mice

Loss of epsins from the endothelium significantly reduced atherosclerotic plaques in the *Apoe*^−/−^ mouse model, as demonstrated by Oil Red O (ORO) staining of the aortic root, arch, and BCA (Brachiocephalic artery) (Supplemental Figure 5), in line with our previous report^29^. Since EndoMT is critical for atherosclerosis, we measured EndoMT markers in aortic roots from *Apoe*^−/−^ and EC-iDKO/*Apoe*^−/−^ mice. As shown in Figure 3A & B, we co-stained CD31 with SMC markers αSMA and Myh11, the overlapping percentile between CD31 and SMC markers is much less in EC-iDKO/*Apoe*^−/−^ mice than in *Apoe*^−/−^ mice, suggesting that loss of endothelial epsins inhibited EndoMT. Similar results were obtained from the co-staining of CD31 with α-SM22 and Calponin in aortic roots (Supplemental Figure 6).

**Figure 3.**
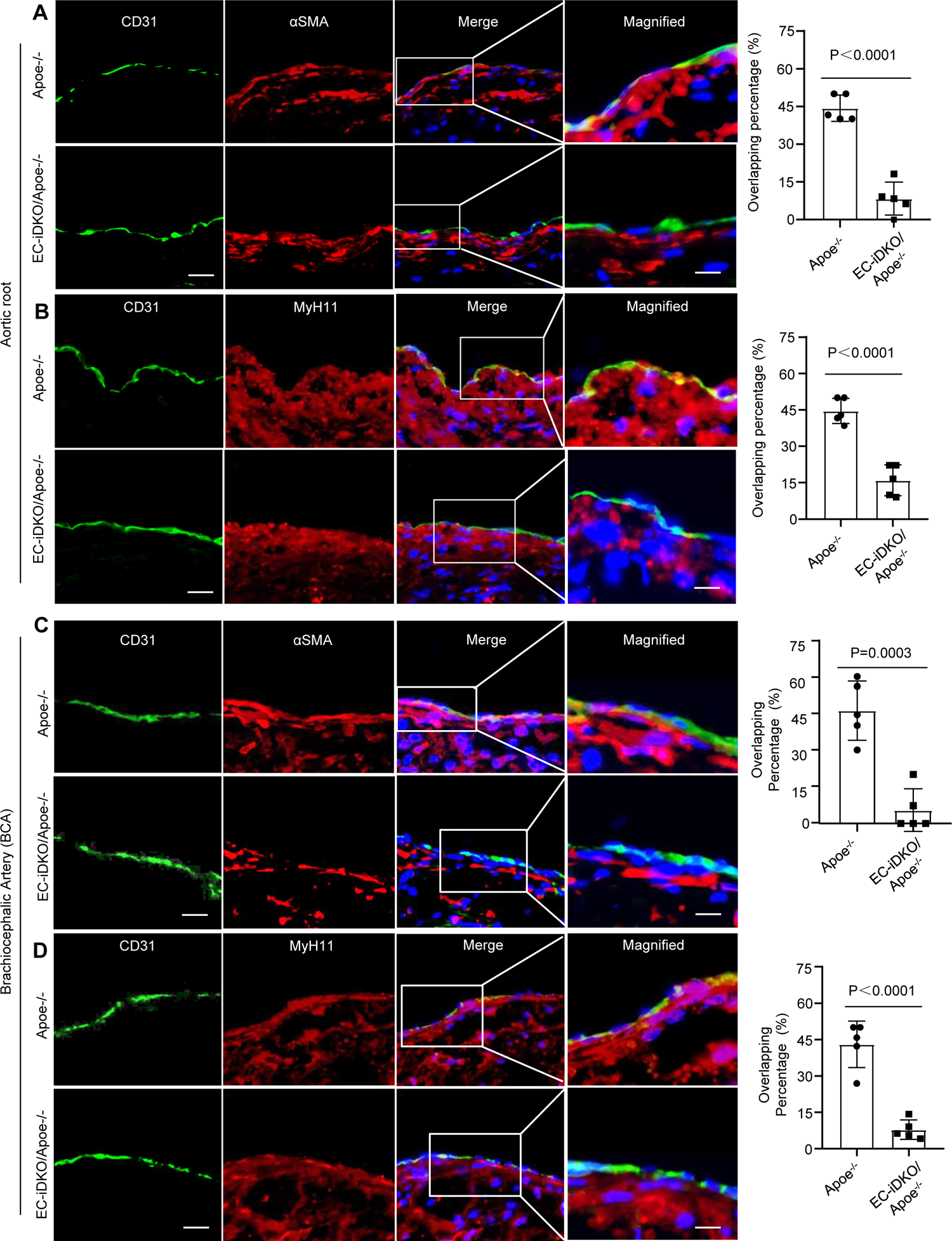
EndoMT is attenuated in EC-iDKO/*Apoe*^−/−^ mice. **A**, **B**, Immunostaining of aortic roots with CD31 and EndoMT markers α-SMA (**A**), MyH11 (**B**) in *Apoe*^−/−^ and EC-iDKO/*Apoe*^−/−^ mice fed WD for 12 weeks. n=5 in each group, P value was indicated in the bar graph. Scale bar: 20 µm, and magnified: 10 µm. **C**, **D**, Immunostaining of BCA with CD31 and EndoMT markers α-SMA (**C**), MyH11 **(D**) were co-stained with CD31 in *Apoe*^−/−^ and EC-iDKO/*Apoe*^−/−^ mice fed WD for 12 weeks. n=5 in each group, P value was indicated in the bar graph. Scale bar: 20 µm, and magnified: 10 µm.

We then measured EndoMT markers in BCA region of arches. As shown in Figure 3C &D, the percentile of CD31 co-staining with SMC markers α-SMA and Myh11 were significantly reduced in EC-iDKO/*Apoe*^−/−^ mice, analogous to the staining for SM22 and Calponin in iDKO/*Apoe*^−/−^ mice (Supplemental Figure 7). Taken together, loss of endothelial epsins inhibits EndoMT in *Apoe*^−/−^ mice.

### Loss of epsins in endothelial cells attenuates the expression of EndoMT markers

Mouse primary cultured aortic endothelial cells (MAECs) were isolated as previously reported^29^. We measured EndoMT markers in MAECs from WT and EC-iDKO after treating with 10 ng/mL TGF-ß for 5 days, and our results demonstrated that VSMC and mesenchymal stem cell markers Acta2, N-cadherin, Collagen 1a and Fibronectin are significantly increased in WT MAECs, but not in EC-iDKO MAECs (Figure 4A, B), while EC cell markers VE-cadherin and Claudin-5 is lower in WT MAECs, but higher in EC-iDKO MAECs in qPCR analysis (Figure 4C; see primers in Supplemental Table 1), further suggesting that epsins promote EndoMT.

**Figure 4.**
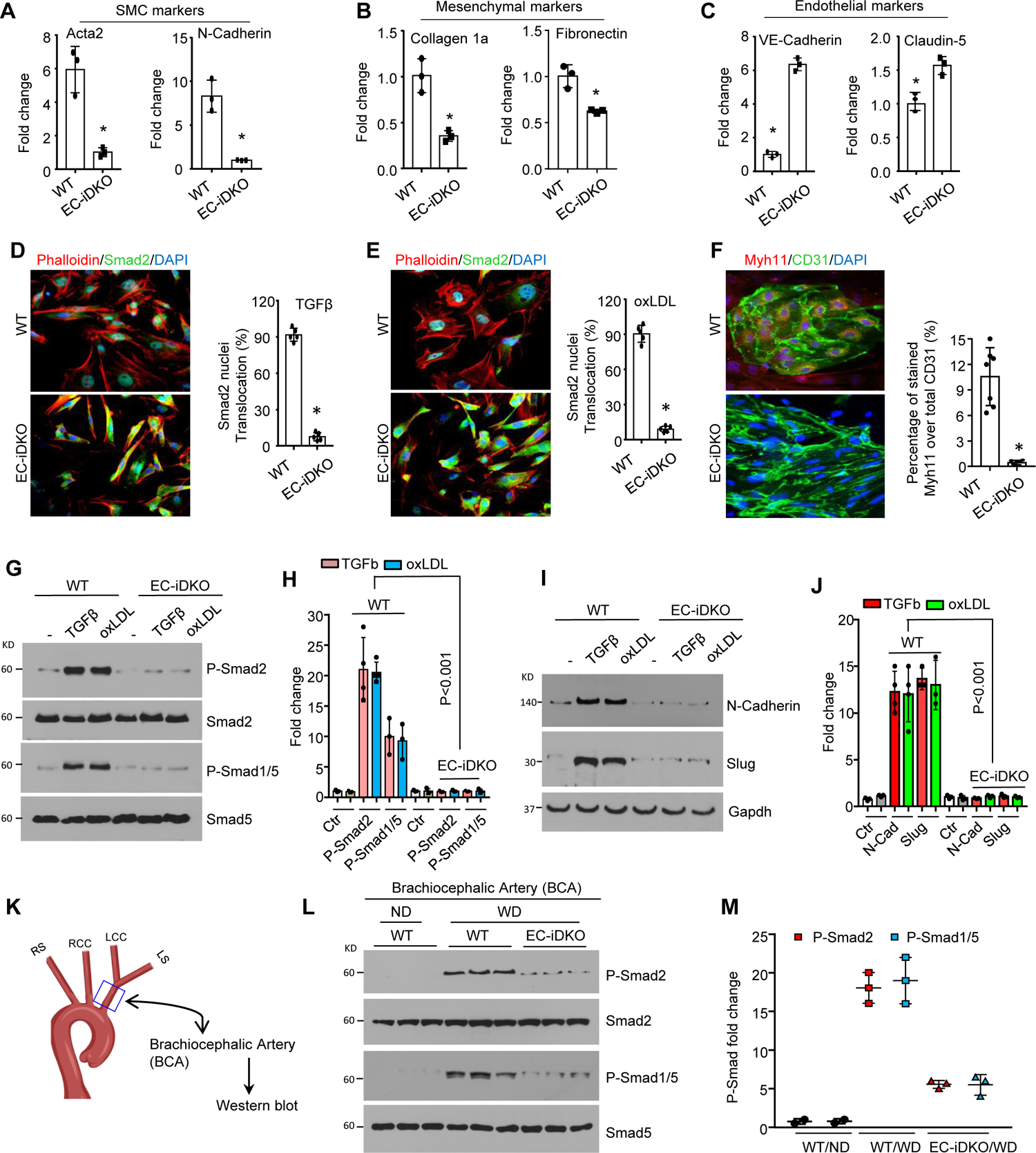
Loss of endothelial epsins inhibits EndoMT via TGF-ß signaling *in vitro* and *in vivo*. **A** to **C**, MAECs of WT and EC-iDKO were treated by 10 ng/mL TGF-ß for 2 days, change fresh medium and TGF-ß every-other-day, followed by qPCR to detect EndoMT markers. SMC markers Acta2, N-cadherin, Mesenchymal markers Collagen 1a, Fibronectin, endothelial markers VE-cadherin and Claudin-5 were measured respectively. n=3, *P<0.01. **D**, **E**, Smad2 translocation was measured by immunofluorescent co-immunostaining Smad2 and Phalloidin-Flu 594. MAECs of WT and EC-iDKO were treated by 10 ng/mL TGF-ß or 100 µg/mL oxLDL for 5 days. n=5, P<0.001. **F**, Co-immunostaining of CD31 and Myh11 for MAECs of WT and EC-iDKO treated by 10 ng/mL TGF-ß for a week. n=7, P<0.001. Scale bars: 50 µm for (**D**, **E**, **F**). Magnified in (**F**): 25 µm **G** to **J**, MAECs of WT and EC-iDKO were treated by 10 ng/mL TGF-ß or 100 µg/mL oxLDL for 5 days, followed by western blot with specific antibodies for the phospho-Smad2 & phospho-Smad1/5 in TGF-ß signaling (**G**, **H**) and target gene expression (**I**, **J**). **K** to **M**, TGF-ß signaling as indicated by phospho-Smad2 and phospho-Smad1/5 was measured in western blot from brachiocephalic artery (BCA) samples of WT and EC-iDKO fed western diet for 8 weeks. WT mice fed normal chow diet (ND) services as basal level controls. (**K**) Cartoon of the aortic region chosen for western blot in brachiocephalic artery. n=3 in each group. * WT/ND *vs* WT/WD, P<0.001; # WT/WD *vs* EC-iDKO/WD, P<0.001.

Functionally, we measured cell migration using Transwell Migration Assay, EC-iDKO cells treated by TGFβ migrated less than WT MAECs inducted in 10% FBS chamber (Supplemental Figure 8).

### Loss of endothelial epsins attenuates TGF-ß signaling *in vitro*

TGF-ß signaling has been proposed to be the main driver of EndoMT through phospho-Smad2 nucleus translocation^8, 11, 12, 14^. To this end, we measured TGF-ß signaling by TGF-ß-induced Smad2 translocation. As shown in Figure 4D & E, Smad2 translocation to nuclei by TGF-ß or oxLDL treatment is drastically inhibited in the EC-iDKO MAECs in immunofluorescent staining (Close-up view provided in the Supplemental Figure 9; also see antibody resources in Supplemental Table 2). To further confirm this result, we co-stained the TGF-ß signaling target gene Myh11 or αSMA with CD31, and we found that approximate 10.5% WT EC cells can be transformed from EC to SMC under TGF-ß treatment for a relatively longer period, while epsin loss in ECs strongly inhibited EndoMT (Figure 4F; Supplemental Figure 9 & 10). Furthermore, we treated MAECs from WT and EC-iDKO mice with TGF-ß or oxLDL. TGF-ß signaling, indicative of phospho-Smad2, phospho-Smad1/5 as suggested by Cooley et al^37^, and the target gene N-cadherin and Slug expression, are significantly reduced in MAECs of EC-iDKO compared to WT-MAECs in western blots (Figure 4G to 4J), in line with the results from immunofluorescence staining (Figure 4D to 4 F).

### Loss of endothelial epsins attenuates TGF-ß signaling *in vivo*

Next, we measured downstream TGF-ß signaling activation in the brachiocephalic artery (BCA) of aortas from WT and EC-iDKO mice fed a WD for 2 months. We found that phospho-Smad2 and phospho-Smad1/5 ^37^, were significantly attenuated in the BCA region of endothelial epsin deficient mice (Figure 4K to 4M), while control mice fed a normal chow diet (ND) had no obvious TGF-ß signaling activation. These results are consistent with our immunostaining of TGF-ß target genes such as αSMA, Myh11, SM22 and Calponin (co-stained with CD31) in aortic roots from *Apoe*^−/−^ and EC-iDKO/*Apoe*^−/−^ mice (Figure 3; Supplemental Figure 6 & 7). Taken together, loss of endothelial epsins inhibited TGF-ß signaling and EndoMT *in vitro* and *in vivo*.

### Loss of epsins inhibits EndoMT through a UIM-dependent Epsin-FGFR1 interaction

FGFR1 is a well-characterized endogenous inhibitor of EndoMT ^8, 11, 12, 14^. Consequently, we measured FGFR1 expression in the presence and absence of epsins. As we show in Fig 5A & B, loss of epsins in MAECs significantly upregulated FGFR1 expression, implying that loss of endothelial epsins physiologically blocks the endocytosis of FGFR1. Importantly, this phenomenon did not change in response to treatment with an inflammatory inducer of TGF-ß (2 ng/mL or 10 ng/mL), suggesting that epsins are required for the endocytosis of FGFR1, regardless of the physiological or pathological conditions. This result was confirmed in human aortic endothelial cells (HAECs) using epsin 1 and 2 siRNA (Figure 5C, D).

**Figure 5.**
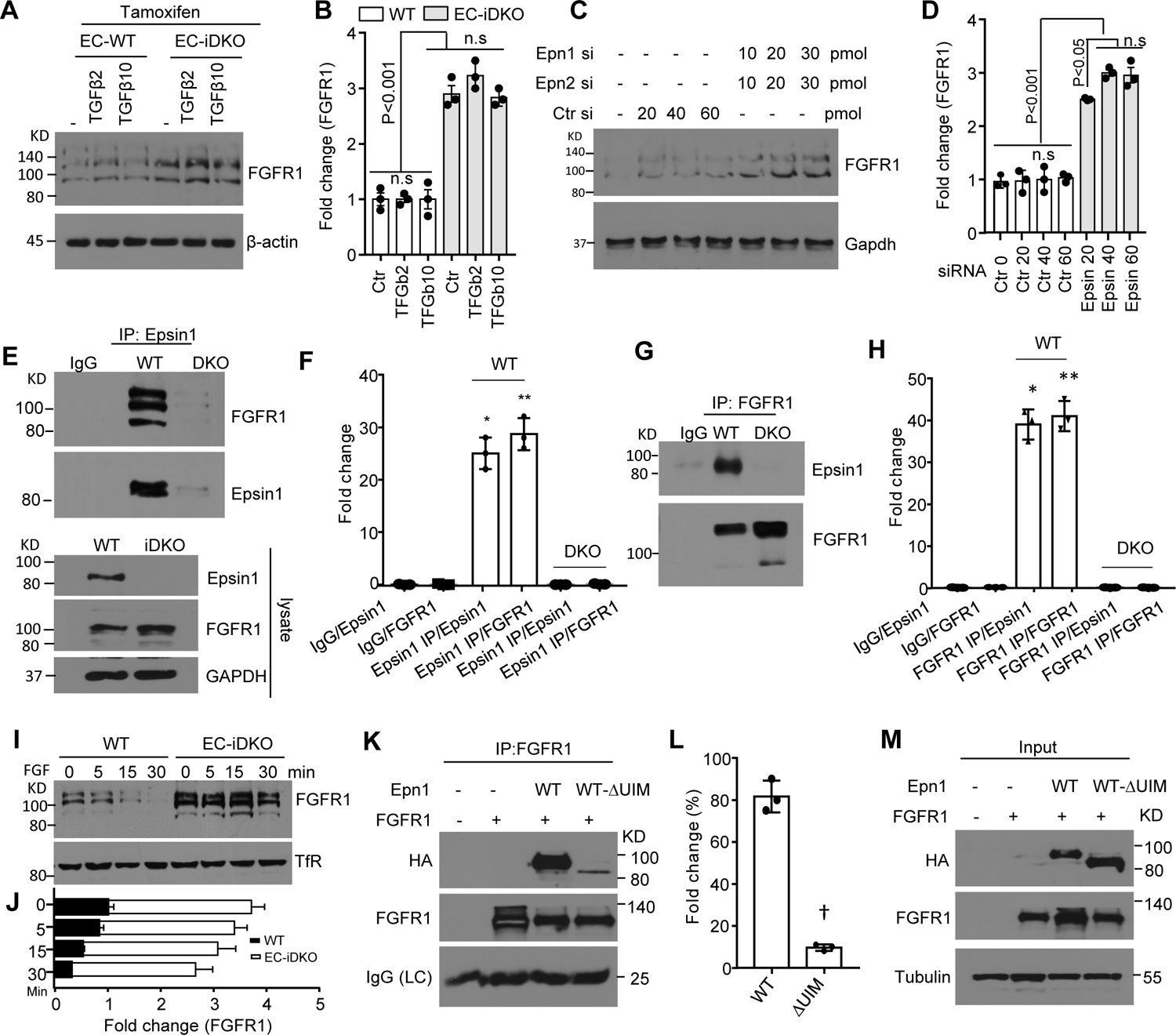
Endothelial epsins promote EndoMT by accelerating UIM-dependent FGFR1 endocytosis and degradation. **A**, **B**, MAECs isolated from WT and Epsin1^f/f^: Epsin2^−/−^:iCDH5-cre mice were both treated with 5 µM tamoxifen for 5 days, followed by the treatment of 2 or 10 ng/mL TGF-ß for 3 days (maintain 1 µM tamoxifen during this stage). Cell lysates were subjected to western blot with specific antibody to FGFR1. n.s, no statistical difference. n=3, * P<0.001. **C**, **D**, Epsin 1 and 2 were knockdown in human aortic endothelial cells (HAECs) by different concentration of siRNA, and cell lysates were subjected to western blot for FGFR1. n=3, * P<0.001. **E** to **H**, Epsin 1 interacts with FGFR1 in reciprocal IP analysis. WT MAECs were cultured in complete EC medium and cell lysates were co-immunoprecipitated with Epsin1 (**E**) or FGFR1 (**G**). IgG as controls in co-IP experiment. Quantification is illustrated in (**F**) and (**H**) respectively. n=3, *, ** P<0.001. **I**, **J**, Biotinylation of PM FGFR1 assay and western blotting for PM FGFR1. **I**, FGFR1 in PM; **J**, decay rate of FGFR1 in WT and EC-iDKO MAECs. n=3, P<0.001. **K** to **M**, HA-tagged Epsin 1 or Epsin 1ΔUIM were co-transfected with FGFR1 plasmid into WT MAECs by electroporation (Lonza), after 36 hours, cell lysis was immunoprecipitated with FGFR1 antibody, followed by western blot with HA antibody. (**M**) Input for the IP analysis in (**K**). **L**, quantification for (**K**), n=3, † P<0.0001.

To evaluate the ability of epsins to directly bind FGFR1, we performed immunoprecipitation analyses. As shown in Figure 5E & F, FGFR1 can be co-immunoprecipitated with epsin 1 in wild type MAECs and *vice versa* (Figure 5G & H). To assess the plasma membrane (PM) FGFR1 levels, we biotinylated PM FGFR1 and precipitated these with a MagnaBind Streptavidin resin (Thermo Scientific). Our results show that loss of endothelial epsins stabilized PM FGFR1 (Figure 5I & J). Furthermore, we co-transfected HA-tagged Epsin1 or HA-tagged Epsin1-ΔUIM and Flag-tagged FGFR1 into WT MAECs by electroporation and immunoprecipitated these proteins with a FGFR1 antibody. We found that deletion of epsin UIM leads to abolition of epsin-FGFR1 interaction (Figure 5K to M; also see endogenous FGFR1 expression in Supplemental Figure 11). In other words, epsin UIM domain is critical for epsin-FGFR1 interaction.

### Strategy to design a synthetic peptide to inhibit EndoMT *in vitro* and *in vivo*

We reasoned that an epsin-UIM-containing peptide would competitively interfere with the bind of epsins to FGFR1 to prevent its endocytosis and degradation (Figure 6A). We synthesized an epsin UIM peptide, which was conjugated to an atheroma-specific targeting peptide sequence Lyp-1^38^ at the N-terminus to form a chimeric peptide, named as API (<u>A</u>theroma UIM <u>P</u>eptide Inhibitor) (Figure 6B). We first tested the peptide penetration capability in MAECs. Cells were pre-treated by 100 µg/mL oxLDL for 6h to create a “pro-inflammation” status, followed by the addition of FITC-conjugated API or control peptides. After 15h, cells were washed and images were taken under fluorescent microscopy. Our result demonstrates that only FITC-API, which contains the atheroma targeting peptide, can efficiently enter cells, but not FITC-UIM peptide (Figure 6C); suggesting that the leading peptide Lyp-1 is indispensable. Pretreatment of MAECs with API peptide considerably inhibited TGF-ß-induced EndoMT compared to control peptide (Figure 6D, E). This result is further supported by Smad2 immunostaining in control or API peptide preloaded MAECs (Figure 6F, G; close-up view provided in Supplemental Figure 12). API peptide pretreatment can significantly block the Smad2 translocation from cytoplasm to nuclei (Figure 6F, G).

**Figure 6.**
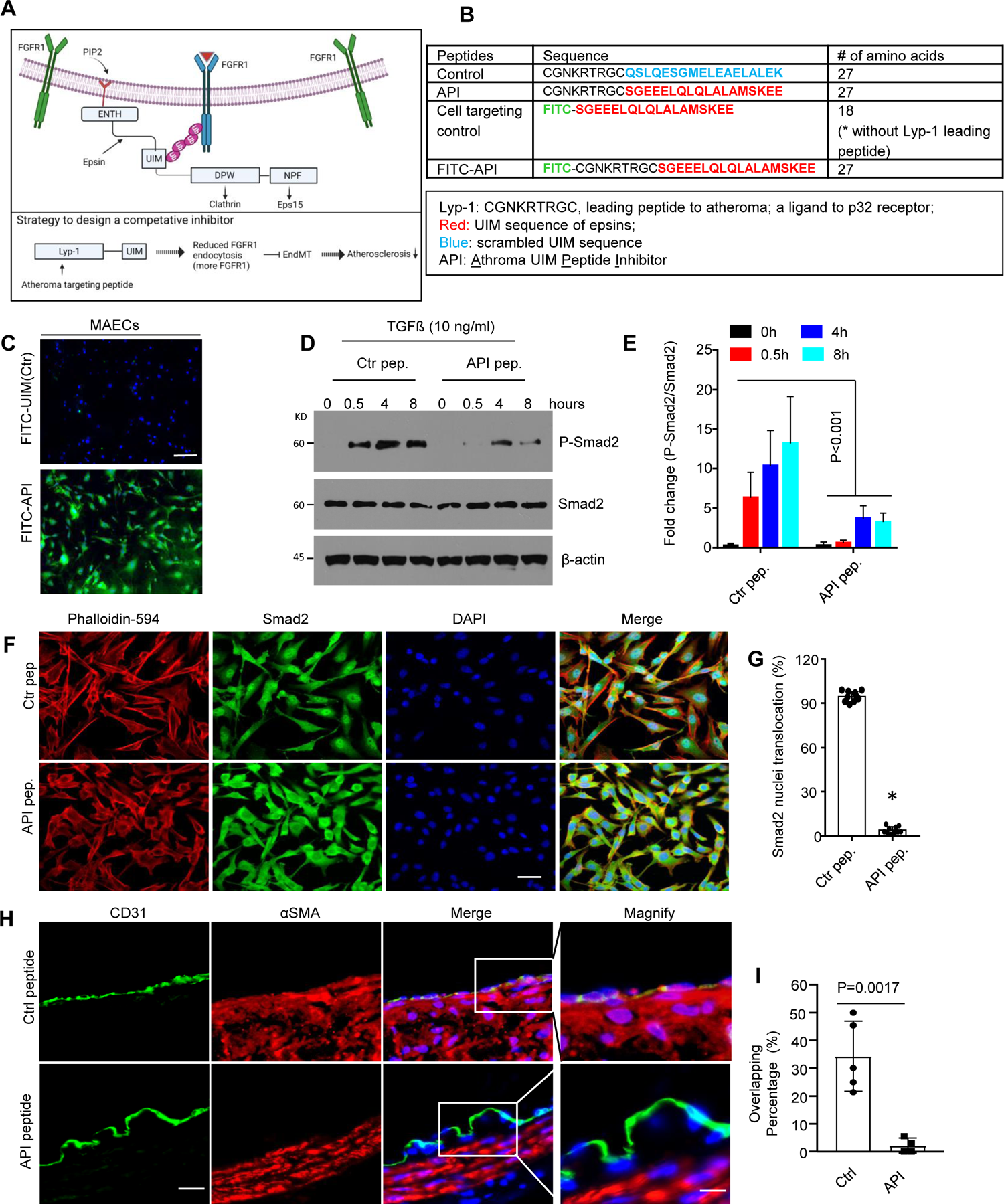
API peptide administration prevents TGF-ß signaling activation *in vitro* and in the *Apoe*^−/−^ model. **A**, Strategy for the design of a synthetic peptide API to competitively bind FGFR1 and block endogenous epsin binding **B**, Peptide sequences used in this study **C**, Co-incubation of 100 µg/mL oxLDL and 25 µM FITC-API or FITC-UIM for 15 h, cells were then fixed and stained with DAPI as reference. Images were captured under Olympus microscopy. Scale bar: 100 µm. **D**, **E**, WT MAECs were pre-loaded with 50 µM control or API peptide, followed by the treatment of TGF-ß (10 ng/mL) as indicated time points. Phospho-Smad2 was measured in western blot and quantified (E). n=3, *P<0.001. **F**, **G**, WT MAECs were pre-loaded with 50 µM control or API peptide, followed by the treatment of TGF-ß (10 ng/mL) for 3 days, and co-stained Smad2 nuclei translocation with Phalloidin-594 (**F**) and quantified (**G**). n=8, *P<0.001. Scale bar: 20 µm. **H**, **I**, Control or API peptide treated mice for 12 weeks on WD was co-stained with CD31 and SMC markers α-SMA (**H**) and quantification (**I**), n=5 in each group, * P<0.01 in α-SMA. Scale bar: 200 µm (F), 20 µm (H).

We then tested the effect of API on EndoMT *in vivo*. We first examined the specific targeting of API peptide by injecting a FITC-conjugated API peptide into *Apoe*^−/−^ mice fed a WD for 6 weeks. Four hours post-injection, mice were sacrificed and aortic roots were stained with DAPI. The API peptide was found to localize to atheroma, especially in the endothelium (Supplemental Figure 13). We compared EndoMT marker αSMA between *Apoe*^−/−^ and EC-iDKO/*Apoe*^−/−^ mice fed WD for 12 weeks. SMC markers αSMA overlapping with CD31 is significantly reduced in API peptide treated group, but rather control peptide group (Figure 6H), suggesting that API block endothelial cells transition to SMC, while FGFR1 is remarkably elevated in API treatment group, which in turn to inhibit TGF-ß signaling (Supplemental Figure 14), in line with our discovery in genetic mouse model, cell culture model and scRNA-seq data.

### Perturbation of Epsin-FGFR1 interaction *in vivo* by API peptide impedes atherosclerotic progression in *Apoe*^−/−^ mouse model: role of prevention

We then performed experiments to test the therapeutic efficacy of API in *Apoe*^−/−^ atherosclerotic mouse model. In *Apoe*^−/−^ atherosclerosis model, mice were injected with API peptide by every-other-day at 50 mg/kg dosage and fed on WD for 2 months. Administration of API significantly inhibited atherosclerotic lesion development of aortic root (Figure 7A, B) and arch (Figure 7C and 7D). To further support our conclusion, we injected API to an alternative carotid atherosclerosis model. Expectedly, API peptide significantly attenuated atherosclerotic lesion development, associated with reduced macrophage infiltration (data not shown). Together, modulation of Epsin-FGFR1 interaction *in vivo* by API peptide is feasible and effective to inhibit EndoMT and atherosclerosis, mimicking the loss of EC epsin phenotype. This result implies a clinical potential to treat atherosclerosis by targeting epsins.

**Figure 7.**
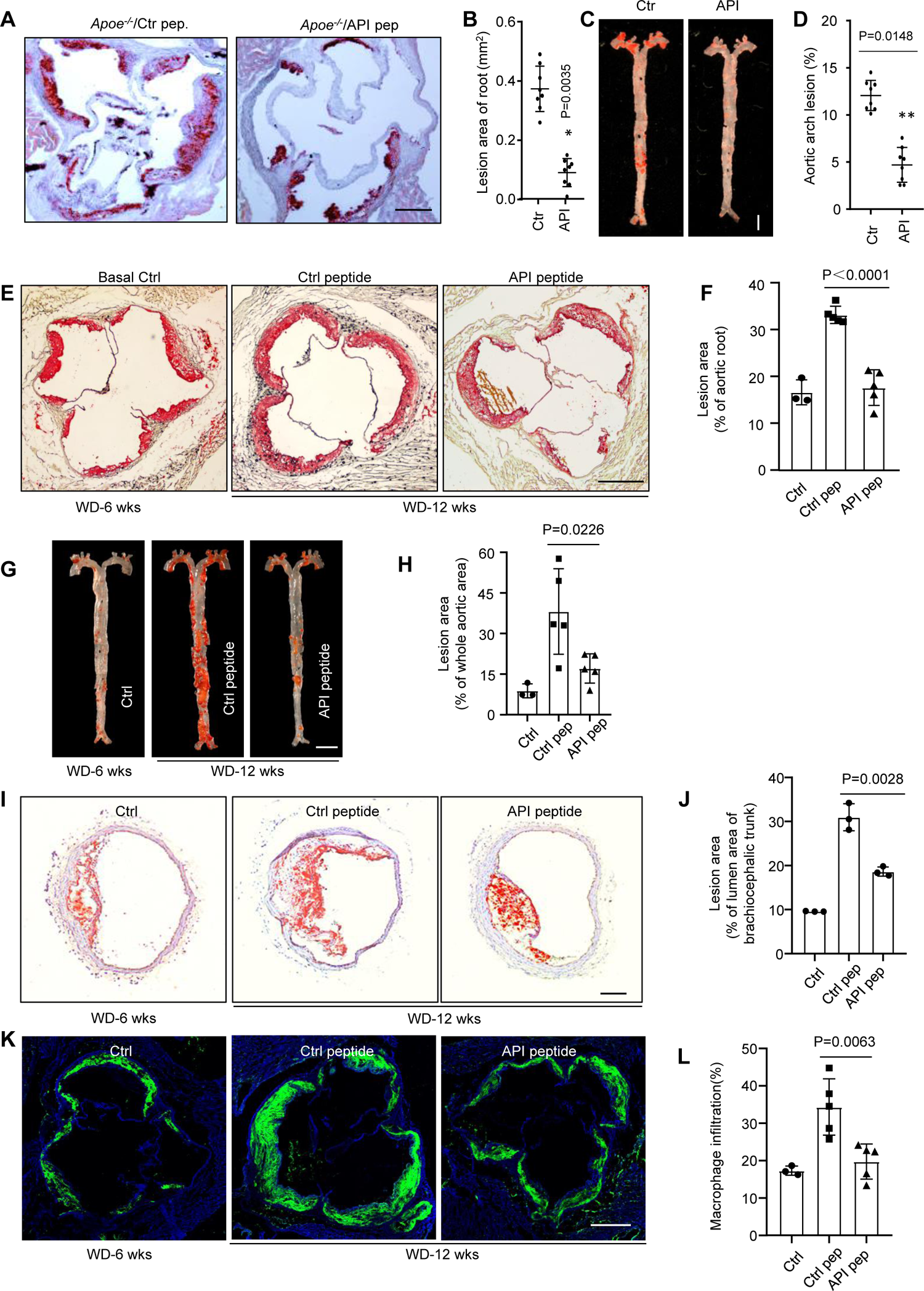
API peptide therapy reduces atherosclerosis in the *Apoe*^−/−^ mouse model. **A** to **D**, Administration of API peptide significantly attenuated atherosclerosis in aortic roots (**A**, **B**) and aortic arches (**C**, **D**) in an *Apoe*^−/−^ animal model. Peptide dose is 25 mg/kg, IV route, and twice a week. n=8 in each group; * or ** P<0.05. Scale bars: 500 µm (**A**); 5 mm (**C**). **E** to **J**, API peptide treatment in lesion-laden *Apoe*^−/−^ models blocked the progression of atherosclerosis. Age matched male *Apoe*^−/−^ mice were fed WD for 6 weeks (sacrificed some mice as basal controls), and started to inject API peptide or control peptide (25 mg/kg, I. V. twice a week) for another 6 weeks, aortic roots (**E**, **F**), arches (**G**, **H**) and BCA (**I**, **J**) were stained with ORO and quantified. Basal control, n=3. Peptide injected group, n=5. P values were labelled in the bar graphs. Scale bar: 500 µm (**E**); 5 mm (**G**); 200 µm (**I**). **K,** Macrophage infiltration staining by CD68 for aortic roots of basal control, control peptide and API peptide treatment groups. n=5, Ctr pep. *vs* pep., P=0.0063. Scale bar: 500 µm (**K**)

### API peptide therapeutic efficacy in the lesion-established *Apoe*^−/−^ mouse model: role of therapy

To determine the therapeutic potential of our candidate peptide compound, we established lesion-laden *Apoe*^−/−^ mouse model by feeding *Apoe*^−/−^ mice for 6 weeks on Western Diet (WD), then we started to inject control or API peptide to these lesion laden *Apoe*^−/−^ model. As we can see in Figure 7E to Figure 7H, atherosclerotic plaques in API peptide treated group were significantly reduced in atherosclerotic lesions in aortic root and arches. A similar result was seen in BCA region of aortic arches (Figure 7I, J) as well as reduced macrophage infiltration detected by CD68 staining (Figure 7K, L). Together, these results suggest that the API peptide can not only prevent atherosclerosis, but also reverse in established atherosclerotic lesions (Figure 7).

In conclusion, we have identified a new molecular mechanism for epsin-modulated atherosclerosis through EndoMT, in which epsins potentiate endothelial-to-mesenchymal (*i.e.,* SMC) transition. A synthetic UIM-containing peptide (API) that interferes with epsin and FGFR1 interaction *in vivo* shows the potential to create a new class of therapeutics for the treatment of atherosclerosis (Figure 8).

**Figure 8.**
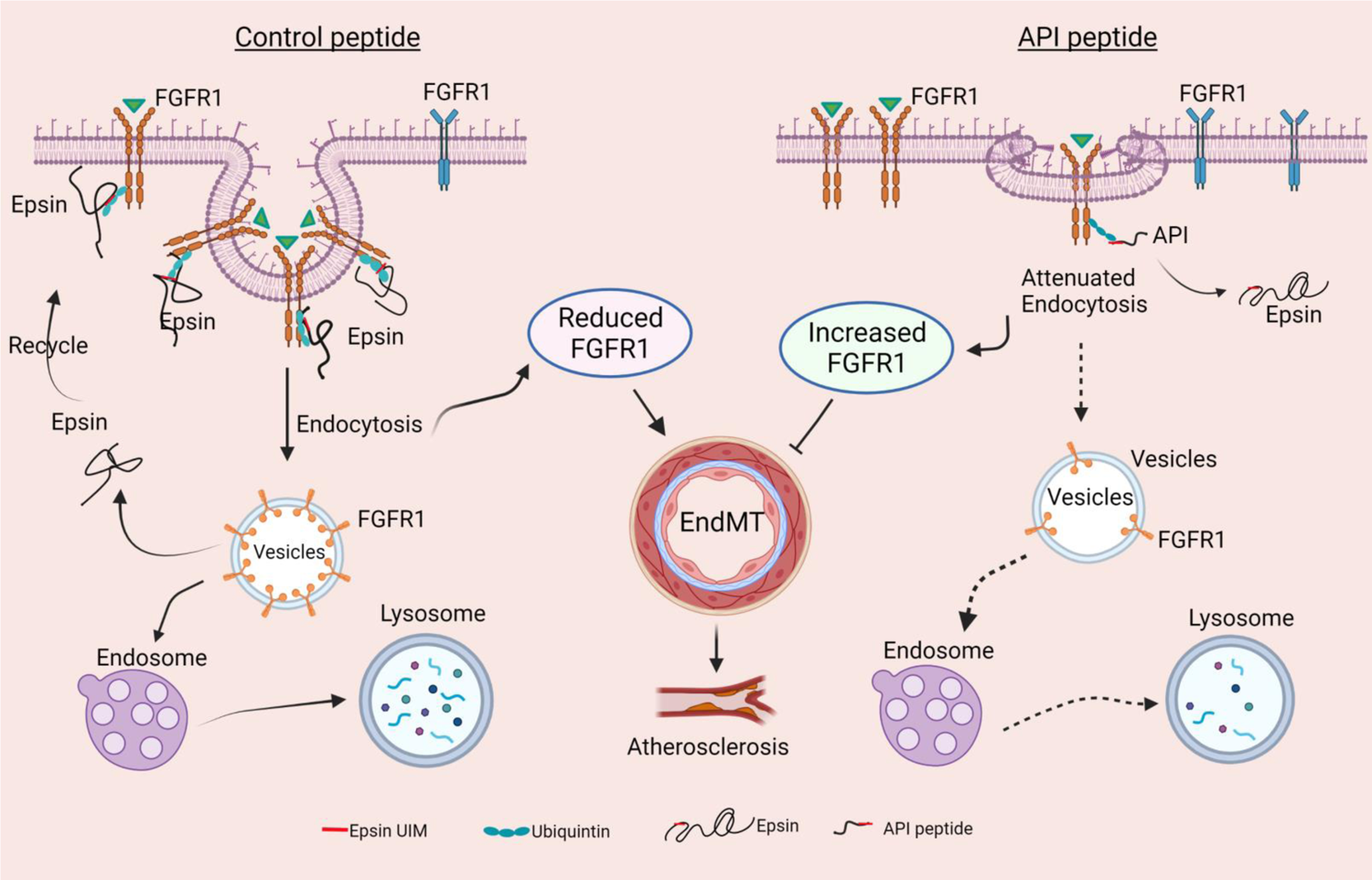
API administration inhibits EndoMT and atherosclerosis in *Apoe*^−/−^ mouse model. Under physiological conditions, Epsins bind FGFR1 via their UIM domain to promote endocytosis, resulting in receptor degradation in the lysosome. Under inflammatory conditions, internalization and degradation of FGFR1 facilitates EndoMT to potentiate atherosclerosis. Application of the synthetic API bind to FGFR1 competitively blocks the ability of endogenous epsins to bind FGFR1, which is replicates genetic epsin deficiency. In both of the latter circumstances, FGFR1 remains at the plasma membrane to inhibit EndoMT induced by inflammation.

## DISCUSSION

Here, we demonstrate a novel role for epsins in potentiating EndoMT using *in vitro* and *in vivo* atherosclerotic models. Using epsin deficient mice, we discovered these endocytic adaptors regulate EndoMT in response to inflammatory stimuli and vascular disease, such as atherosclerosis^6^.

Using scRNA-seq, epsin mutant mice, *Apoe*^−/−^ model, lineage tracing mice and *in vitro* cell culture approaches, we demonstrate that epsins promote EndoMT (Figure 1-4). Mechanistically, epsins modulate the protein levels of FGFR1 (Figure 5)—a well-documented endogenous inhibitor of EndoMT^8, 11, 12, 14^. Loss of endothelial epsins causes the accumulation of FGFR1 on the plasma membrane, which inhibits EndoMT under inflammatory conditions. This discovery expanded our understanding about the mechanism behind EndoMT and the complexity of atherosclerosis. Furthermore, epsins bind FGFR1 (Figure 5), which provides the therapeutic foundation to modulate EndoMT and atherosclerosis.

FGFR1 binds epsins via their UIM to enable receptor endocytosis in physiologically and pathologically circumstances (Figure 5). Conversely, the loss of endothelial epsins stabilizes cell surface FGFR1 (Figure 5), which is crucial for suppression of EndoMT. This finding is similar to our observations for VEGFR2 in neo-angiogenesis and tumor angiogenesis, both of which are dependent on UIM interactions between epsins and VEGF receptors^39, 40^. Interestingly, VEGFR2 has been reported to inhibit EndoMT^41^, however, in large arteries like the aorta, VEGFR2 expression is minimal^23^. Therefore, the upregulation of FGFR1 in large vessels resulting from endothelial epsin loss may suppress EndoMT.

We previously reported that epsins bind to numerous receptors (*e.g.,* IP3R1, TNFR1, TLR2/4, and LRP1) via the UIM domain^30, 42–44^. This suggests that epsin UIM was a critical domain to regulate inflammatory receptors, which was the basis for the development of our therapeutic peptide. As a result, we designed an atheroma-targeting peptide called API, in which a Lyp-1 transduction peptide was fused to the N-terminal of UIM sequence (Figure 6B).

Lyp-1 has been suggested to home to macrophage and p32 positive tumor cells, as well as tumor lymphatic vessels^38, 45, 46^. In our *in vitro* experiments, we found that a FITC-conjugated API peptide was efficiently taken up by MAECs pre-treated with oxLDL. In *Apoe*^−/−^ mice, we visualized the FITC-conjugated API peptide homing to the atheroma (predominantly in the EC layer) (Supplemental Figure 13). In our proof-of-concept study, we provided evidence that atherosclerosis can be dramatically alleviated by downregulating EndoMT and upregulating FGFR1 expression using this atheroma targeted peptide (Supplemental Figure 14). Our data showed that atherosclerotic plaques can be significantly reduced by API peptide administration (Figure 7). However, we could not completely rule out the possibility that some of the API peptide targeted macrophages, which could provide additional anti-inflammatory benefits^30, 42–44^.

Although EndoMT has been implicated in contributing to cardiovascular diseases, including atherosclerosis, pulmonary hypertension, tissue fibrosis and brain vascular malformation^6, 11, 47^, a recent report suggests that EndoMT can transiently stabilize the atherosclerotic fibrous cap^9^, reflecting that the role of EndoMT in atherosclerosis could be dual. The report showed convincingly that in murine and human lesions, 20% to 40% of the protective ACTA2+ myofibroblast (MF)-like cells in the fibrous cap, are actually derived from non-SMC sources, including endothelial cells (ECs) or macrophages that have undergone an endothelial-to-mesenchymal transition (EndoMT) or a macrophage-to-mesenchymal transition (MMT), respectively, suggesting that EndoMT is “protective” to plaque stabilization. However, in more advanced lesions, the beneficial role of EndoMT in plaque stabilization is diminished as PDGFβ signaling-mediated SMC recruitment to the fibrous cap is imperative to reinforce plaque stability. Thus, it is conceivable that EndoMT under aforesaid conditions would further fuel endothelial activation and dysfunction owing to prolonged loss of pool of endothelial cells that may be required for endothelial repair, crucial for maintaining endothelial homeostasis. Nevertheless, the role of endothelial epsins played in EndoMT on the stabilization of fibrous cap of atherosclerosis warrants further investigation. Besides, EndoMT is required for heart mitral valve development and it is detectable for healthy adult valves in sheep and humans^48^. EndoMT can also be reactivated in the postnatal setting in both physiologically and pathologically states^49^. EndoMT has been proposed to be harmful to heart repair^50^, although it is still controversial if fibroblasts derived from EndoMT are responsible for the aforesaid events as recent studies show that a local pool of fibroblasts get reactivated and dynamically remodel following myocardial infarction injury, and lose SMA expression to maintain heart structure post myocardial infarction^51^.

In summary, we demonstrated that the loss of epsins in endothelial cells inhibits atherosclerosis, at least in part, through inhibiting EndoMT. Mechanistically, epsins modulate FGFR1 expression and its endocytosis in atherosclerosis. Targeting atheroma-specific epsins by UIM-containing peptide API mitigates EndoMT and atherosclerosis (Figure 8). These findings are perspective to control atherosclerosis.

## CLINICAL PERSPECTIVE

### What Is New?

* Endothelial epsins are required for EndoMT *in vitro* and *in vivo*.

* Epsins interact with FGFR1 in a UIM-dependent fashion to govern FGFR1 endocytosis and degradation.

* Targeted delivery of a UIM-containing peptide (API) reduces atherosclerotic lesion size.

### What Are the Clinical Implications?

* Targeting epsin-FGFR1 interaction with a therapeutic peptide is feasible means to reduce atherosclerotic plaque burden in mice.

* The homology of the UIM between humans and mice indicates this form of therapy could be used to treat atherosclerosis in patients.

## ACKNOWLEDGEMENTS

Y.D., B.W., and H.C. designed the experiments, analyzed and interpreted the data. Y.D. and B.W performed most of the experiments. M.D. analyzed the scRNA-seq data. B.Z. did the barcoding and cDNA library construction. K.C, S.B., and S.D. helped with immunostaining. S.W. performed the mouse genotyping and colony maintenance. Y.D., D.B.C., and H.C. wrote the manuscript. BGI-Hong Kong performed the scRNA-sequencing. M.L. and K.C. made constructive suggestions. We thank Dr. Yanqiang Li for his preliminary analysis of scRNA-seq data.

## SOURCES OF FUNDING

This work was supported in part by the National Institute of Health (grants R01HL0093242-12, R01HL137229-04, R01HL1563626-01, R01HL158097-01, and R01HL146134-03 to H.C.).

## DISCLOSURES

None

## Supplemental Figures and Tables

**Supplemental Figure 1.**
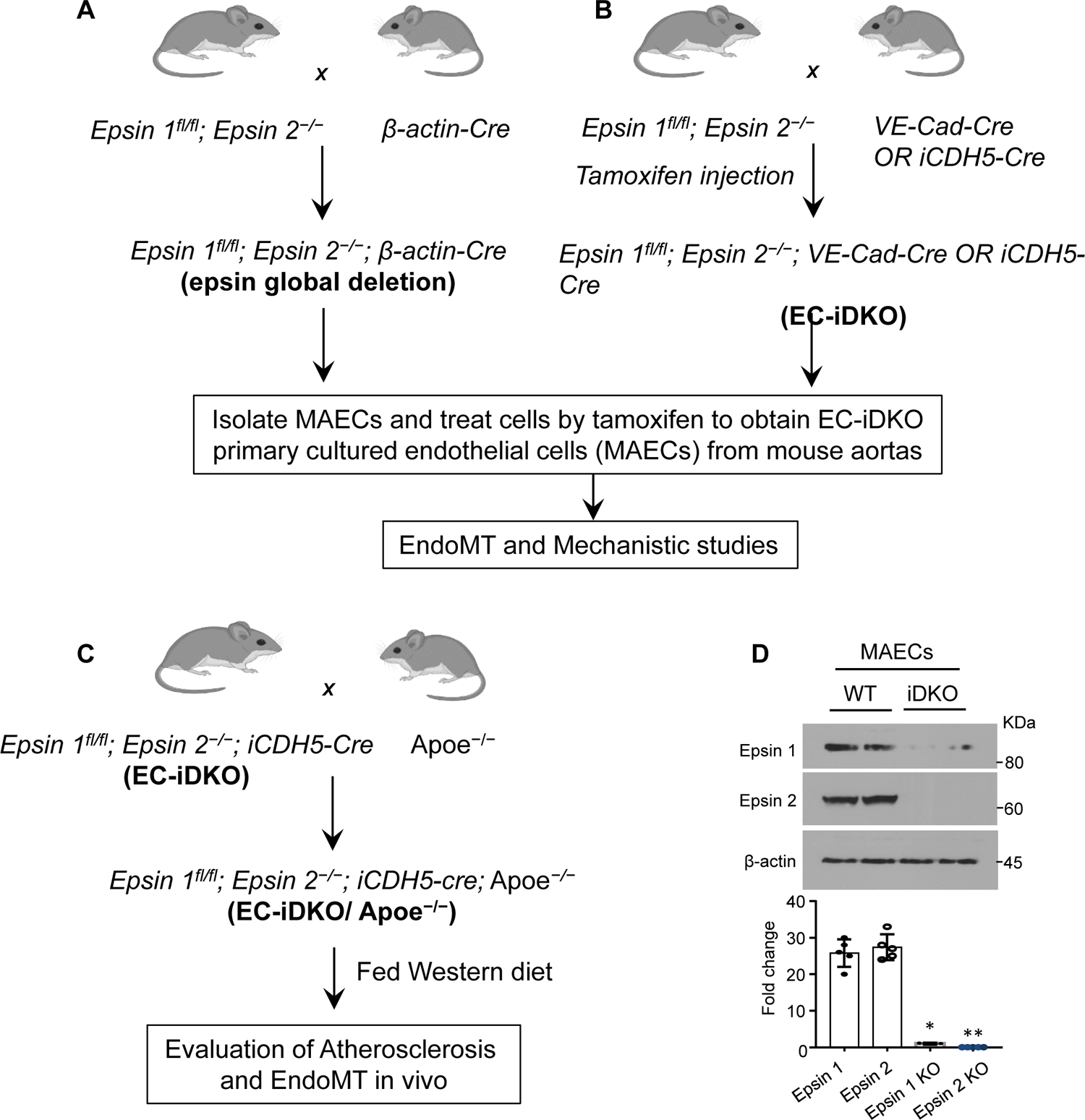
Procedures to generate primary cultured mouse endothelial cells (MAECs)from mouse aortas and epsin mutant mice in Apoe-null murine atherosclerosis model. (**A**) Global epsin DKO under constitutive β-actin driven Cre. (**B**) conditional DKO epsins in endothelium Driven by VE-Cadherin promoter. (**C**) generating epsin DKO mice in Apoe-null mouse model. (**D**) creation of epsins DKO MAECs in petri dishes. Isolated cells were treated by 5 µM tamoxifen for 4 days, followed by western blot analysis. VE: vascular endothelium. n=3. *, ** P<0.0001.

**Supplemental Figure 2.**
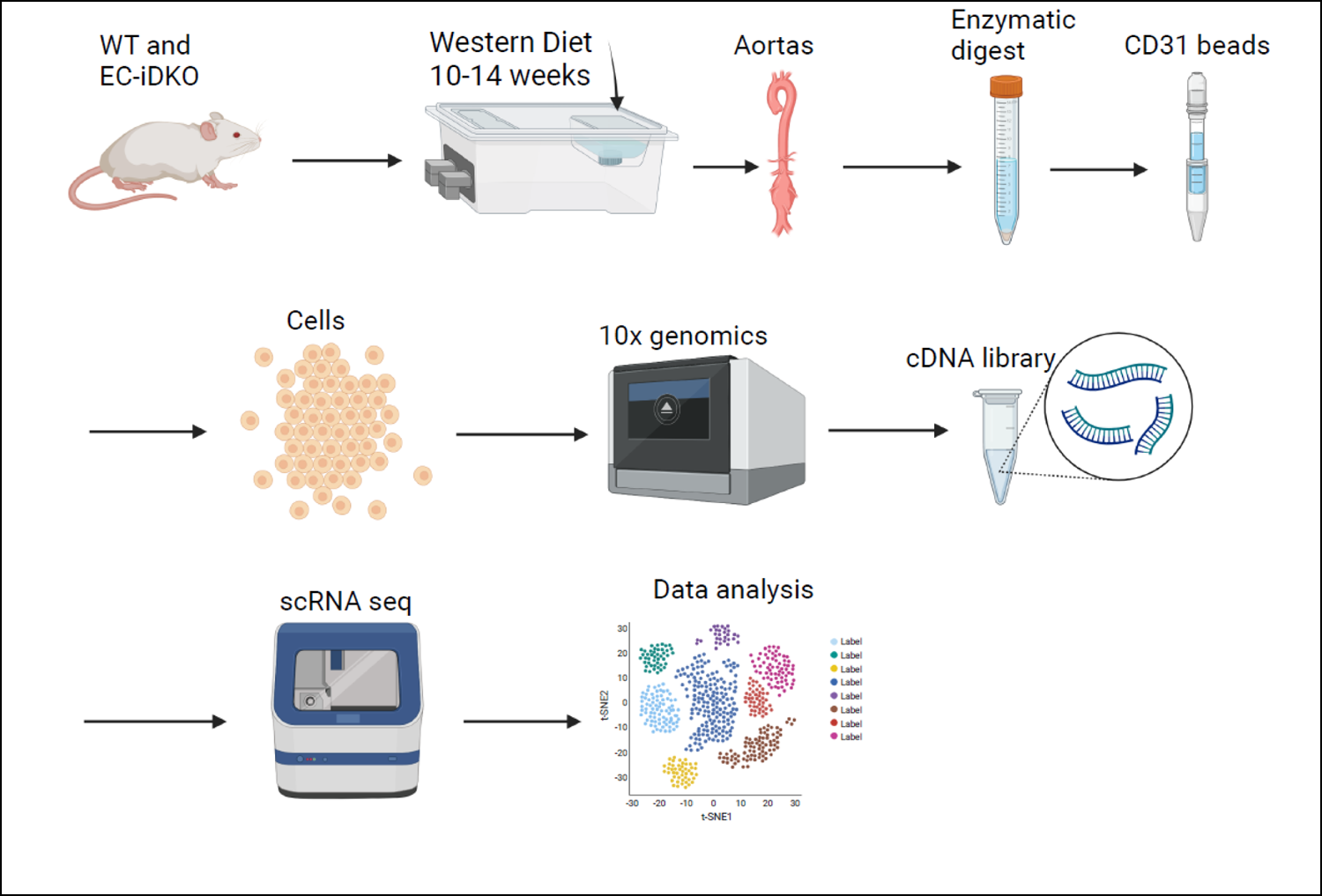
Procedure for single cell RNA sequencing (scRNA seq). WT and EC-iDKO mice were fed WD as indicated time, followed by the isolation of aortas, enzyme digestion and pass through CD31 beads to enrich endothelial cells, and the barcoding cells in 10 x genomics as company instruction. cDNA library was constructed and then perform scRNA seq and data analysis.

**Supplemental Figure 3.**
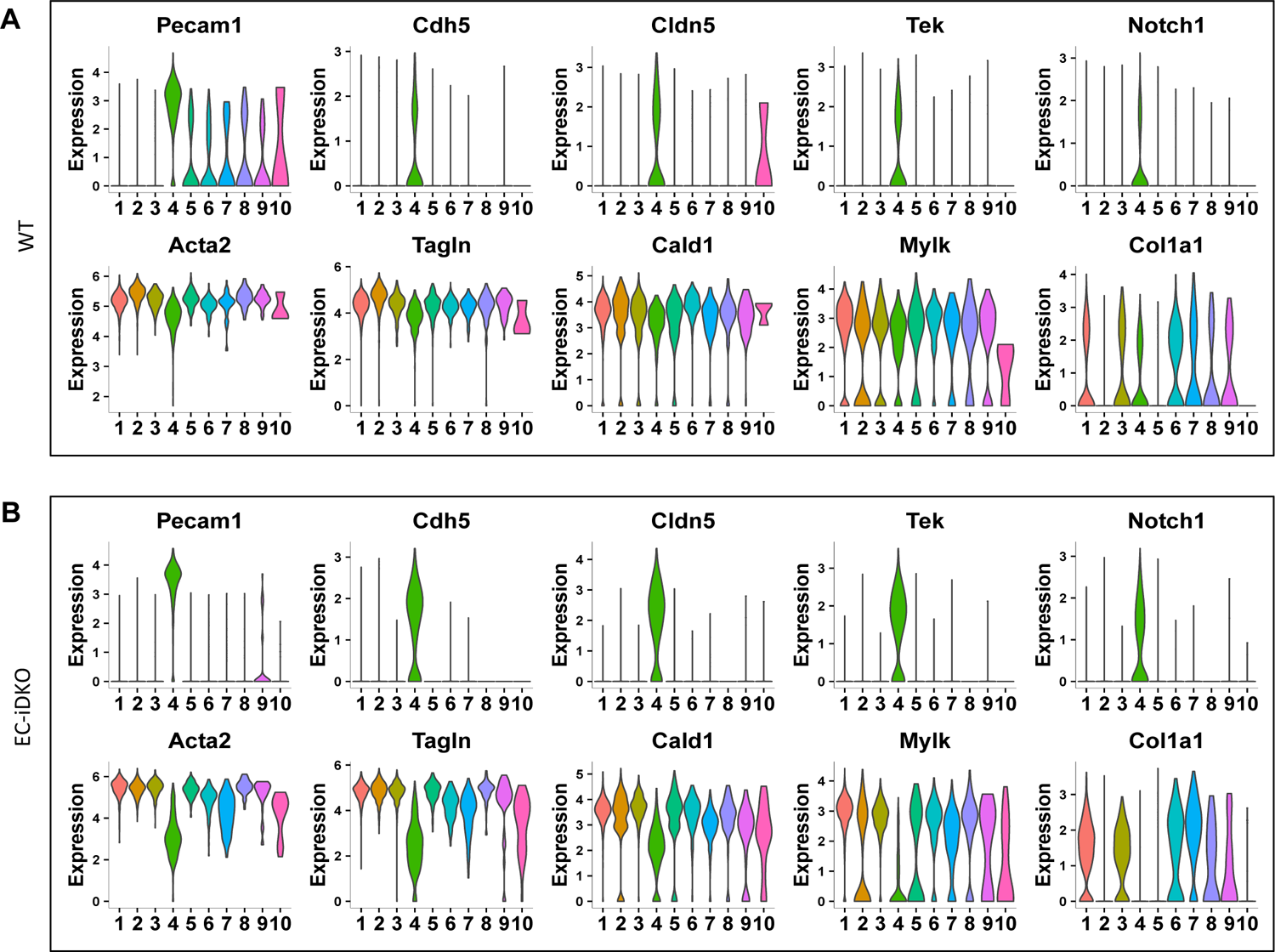
Clustering annotation for single cell transcriptome in EndoMT models. Violin plot of EC and SMC markers in EC-iDKO and WT cells. EC markers: Pecam1, Cdh5, Cldn5, Tek and Notch1; SMC markers: Acta2, Tagln, Cald1, Mylk and Col1a1. (**A**) WT; (**B**) EC-iDKO

**Supplemental Figure 4.**
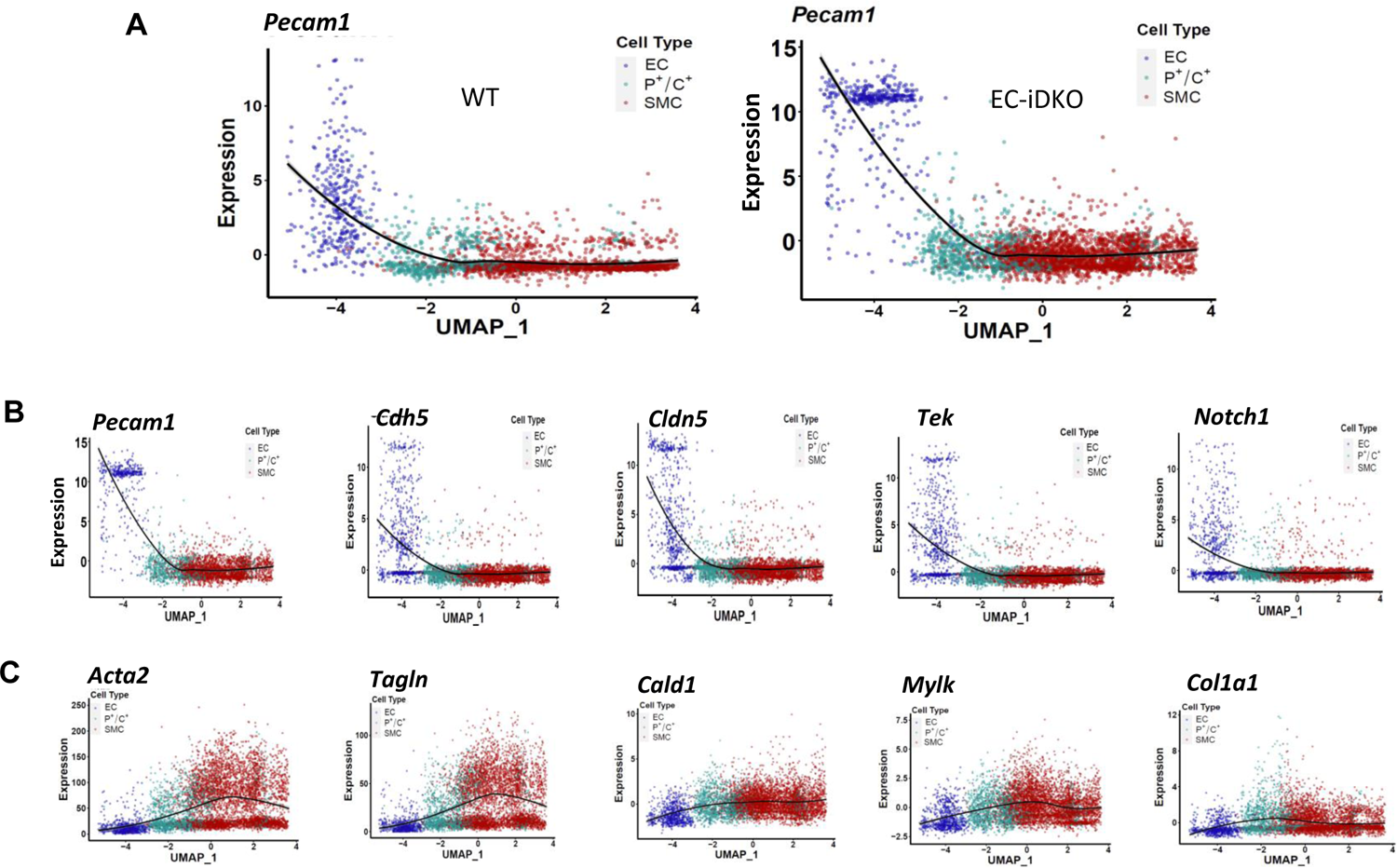
Trajectory plots suggest EndoMT in the integrated cell population. (**A**) Pecam1 plot in EC-iDKO and WT cell population; **(B**, **C**) Trajectory Plot of EC markers and SMC markers in integrated cell population.

**Supplemental Figure 5.**
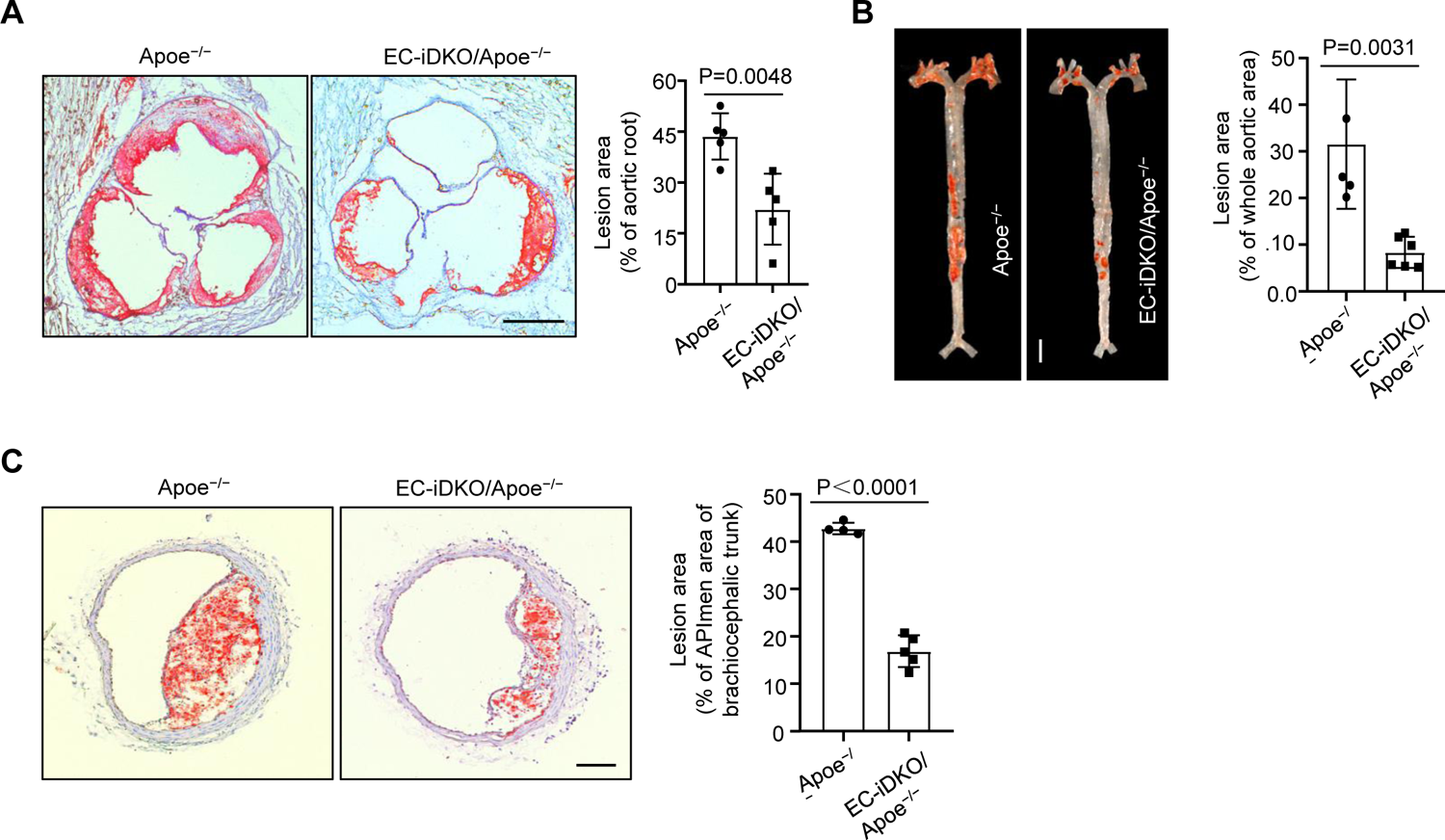
Loss of endothelial epsins reduces atherosclerosis in Apoe^−/−^ atherosclerotic models. (**A**, **B**) ORO staining for aortic roots (**A**) and arch (**B**) of Apoe^−/−^ and EC-iDKO/Apoe^−/−^ mice fed WD for 12 weeks. n=5 in each group, p=0.0031. Scale bar: 500 µm (**A**); 5 mm (**B**). (**C**) ORO staining for BCA from the mice as above, n=5 in each group, P<0.0001. Scale bar: 200 µm.

**Supplemental Figure 6.**
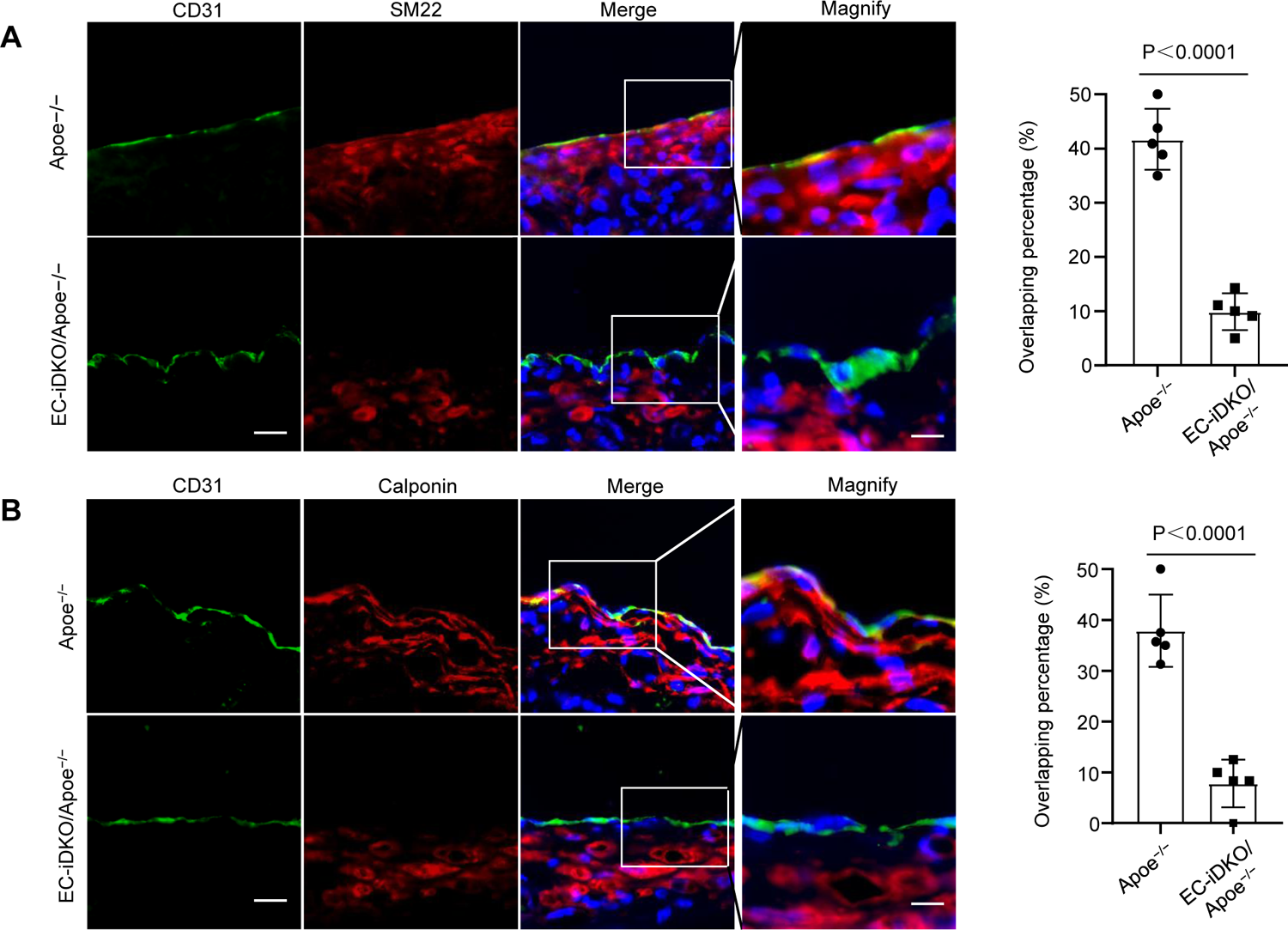
Loss of endothelial epsins inhibits EndoMT in aortic root. Apoe^−/−^ or Epsin1f/f: Epsin 2^−/−^: iCDH5-cre was injected tamoxifen to delete Epsin1 in vivo, and mice were fed WD for 12 weeks. (**A**) CD31 and SM22 co-staining. n=5 mice in each group, P<0.0001. (**B**) CD31 and Calponin co-staining. n=5 mice in each group, P<0.0001. Scale bars: E& F, 20 µm, Magnified, 10 µm.

**Supplemental Figure 7.**
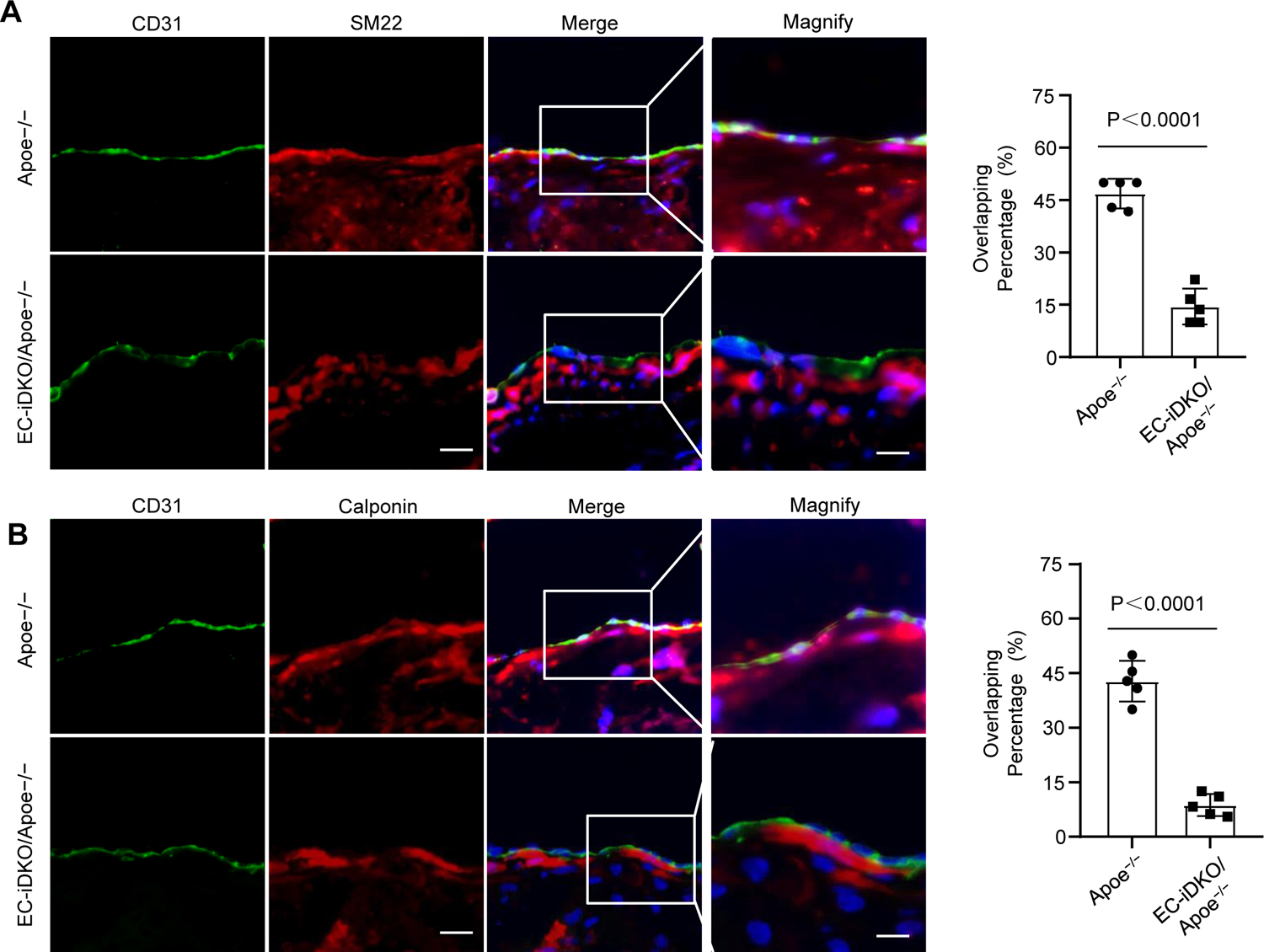
Loss of endothelial Epsins inhibits End MT markers in the BCA region of aortas of Apoe^−/−^ atherosclerotic model. (**A**) CD31 and SM 22 co-staining. n=5 mice in each group, P<0.0001. (**B**) CD31 and Calponin co-staining. n=5 mice in each group, P<0.0001. Scale bars: A& B, 20 µm, Magnified, 10 µm.

**Supplemental Figure 8.**
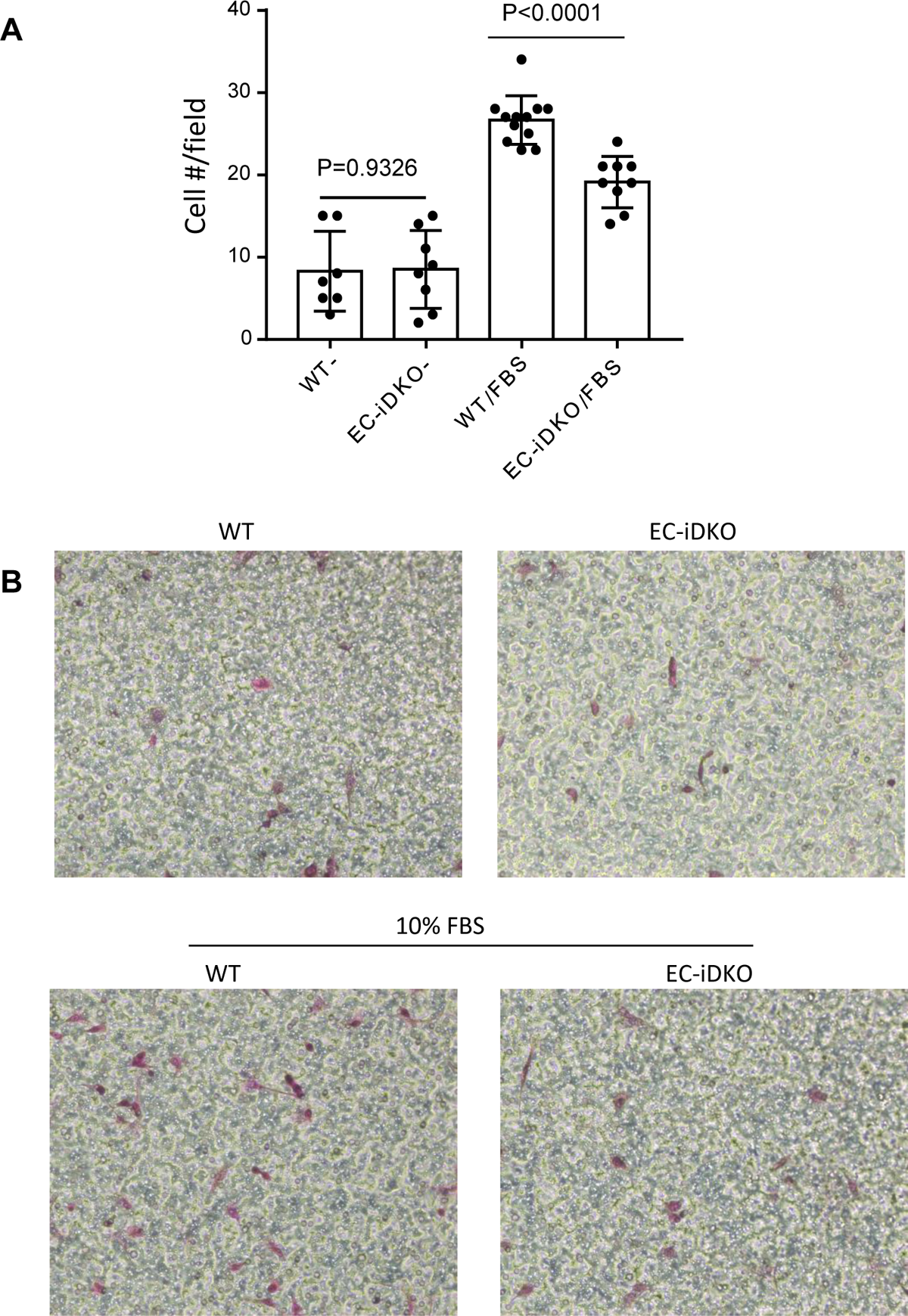
Migration assay of MAECs of WT and iDKO treated by TGFβ1. MAEC cells treated by 10 ng/ml TGFβ for 7 days, followed by Transwell migration assay (7 hours). **(A)** Statistical result from Transwell Migration Assay; n=5 field in three independent experiments; **(B)** Representative images of WT and EC-iDKO cells treated by TGFβ and migration assay after 7 hours in Transwell.

**Supplemental Figure 9.**
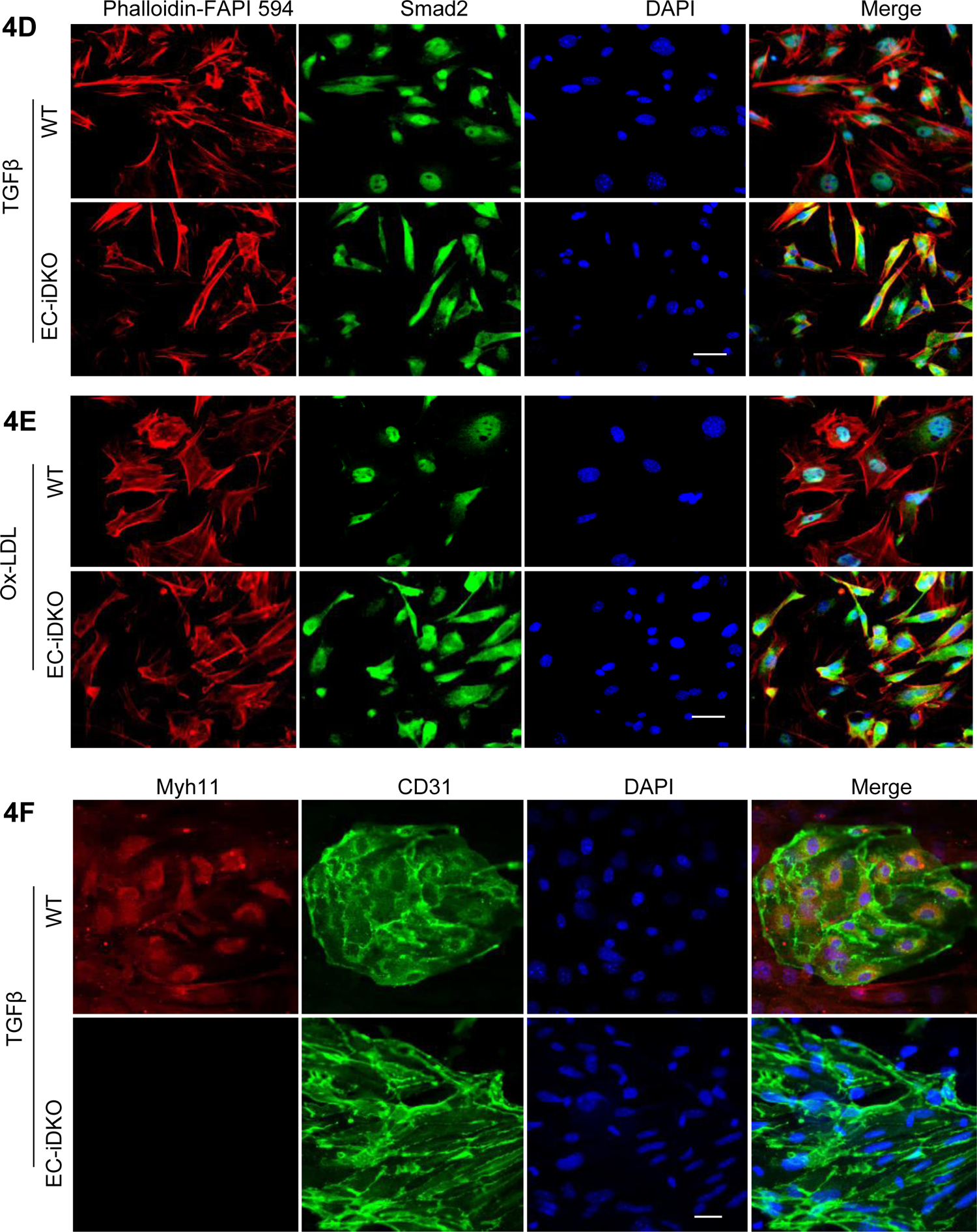
Close-up view (individual panels) of Figure 4D, 4E and 4F. 4D, 4E, Smad2 translocation was measured by immunofluorescent co-immunostaining Smad2 and Phalloidin-Flu 594. MAECs of WT and EC-iDKO were treated by 10 ng/ml TGFβ or 100 µg/ml oxLDL for 5 days. n=5, P<0.001. 4F, Co-immunostaining of CD31 and Myh11 for MAECs of WT and EC-iDKO treated by 10 ng/ml TGFβ for a week. n=7, P<0.001. Scale bars: 50 µm for (D, E, F). Magnified in (F): 25 µm

**Supplemental Figure 10.**
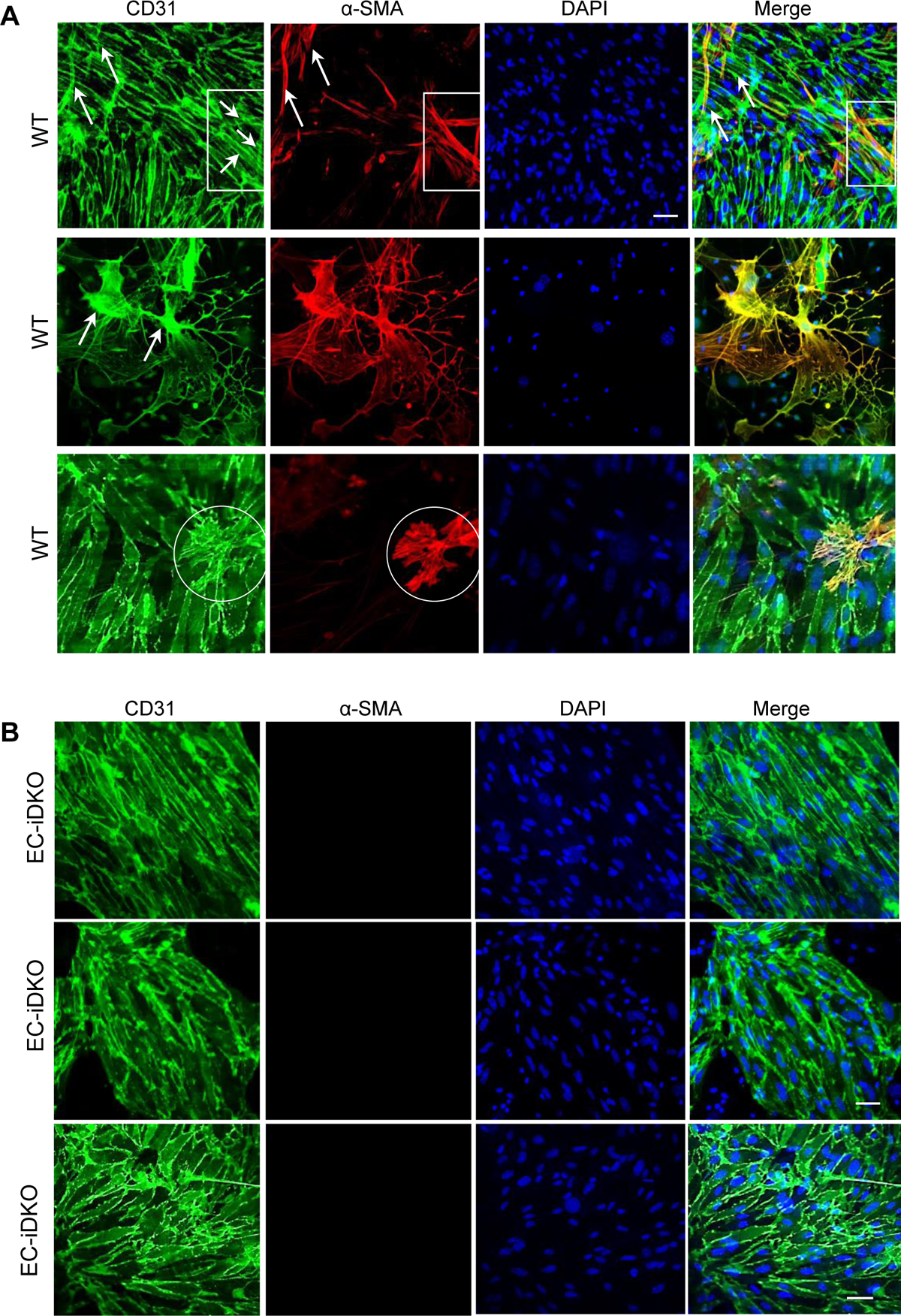
Loss of epsins in endothelium attenuates EndoMT markers promoted by TGFß treatment in vitro, showing more representative EndoMT examples in WT MAECs. MAECs isolated from WT or EC-iDKO mice and treated by TGFβ (10 ng/ml) for 15 days. TGFβ will be replaced every-other-day. During the treatment time, tamoxifen will be always supplied in the medium at 1 µM for both WT and EC-iDKO cells. Circle: SMC-like cells; Arrows: EC is becoming SMC-like cells induced by TGFβ. Scale bar: 25 µm.

**Supplemental Figure 11.**
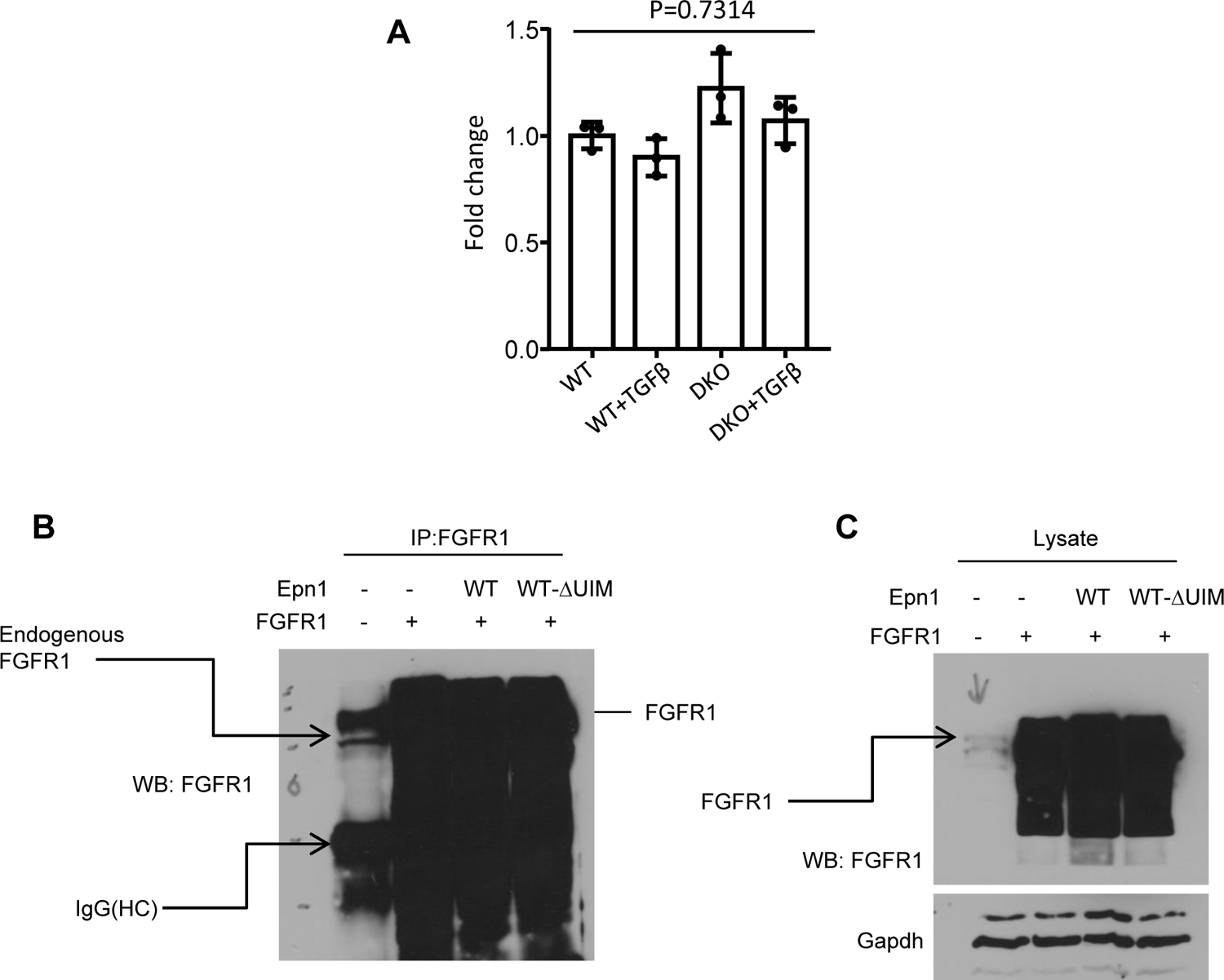
mRNA expression of FGFR1 in MAECs of WT and DKO and Longer time exposure of western blot in main Figure 4K and 4M. (**A**) qPCR test for FGFR1 transcriptional expression in MAECs of WT and DKO cells with or without TGFβ (10 ng/mL) treatment for 24 hours; (**B**, **C**) Longer time exposure of western blot in main Figure 4K and 4M, i.e. IP and input samples, showing endogenous FGFR1 bands in IP (Fig. 4K) or cell lysate (Fig. 4M) separately.

**Supplemental Figure 12.**
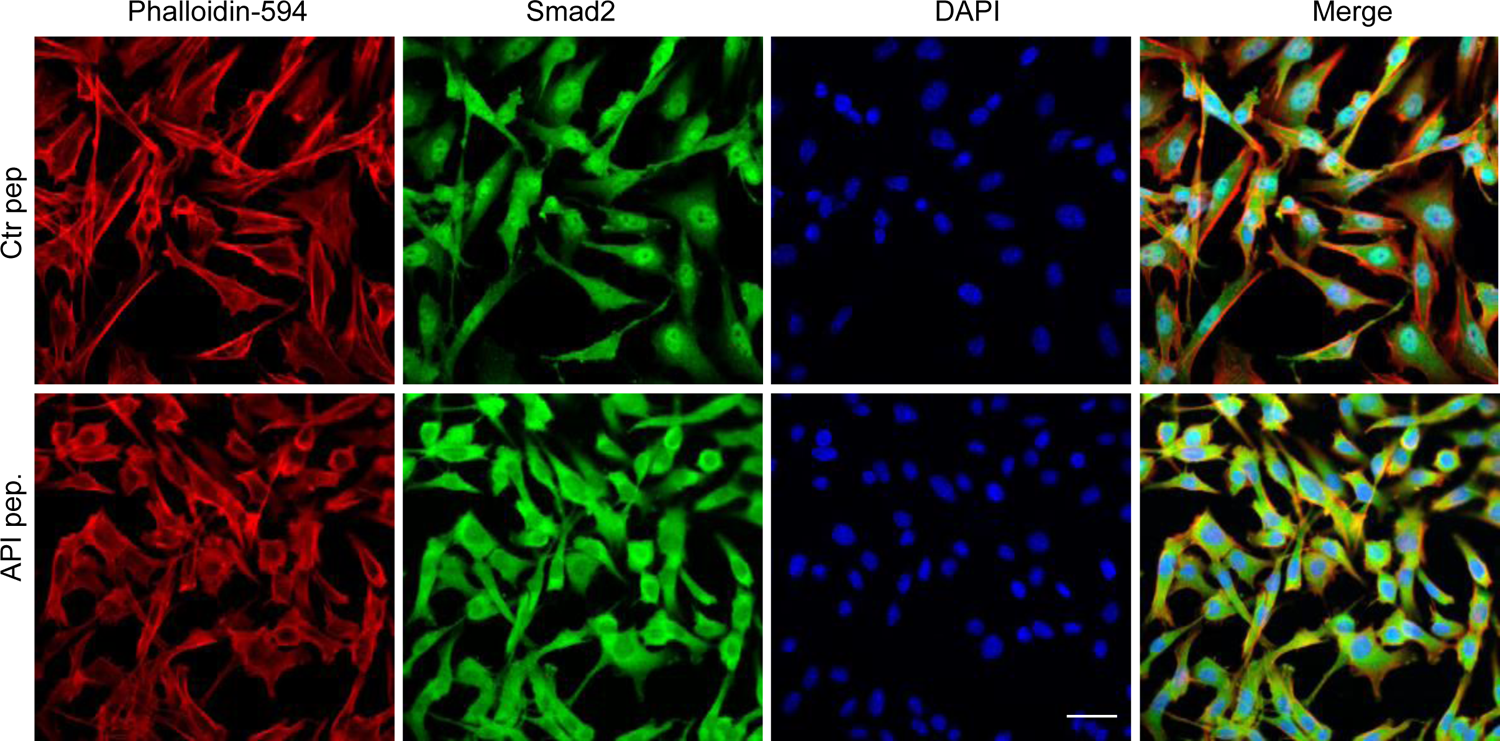
Close-up view (enlarged view) of Figure 6F. Showing Smad2 nuclei staining in Ctr peptide, and API peptide inhibits nuclei translocation. WT MAECs were pre-loaded with 50 µM control or API peptide, followed by the treatment of TGFβ (10 ng/ml) for 3 days, and co-stained Smad2 nuclei translocation with Phalloidin-594; n=8, *P<0.001. Scale bar: 20 µm.

**Supplemental Figure 13.**
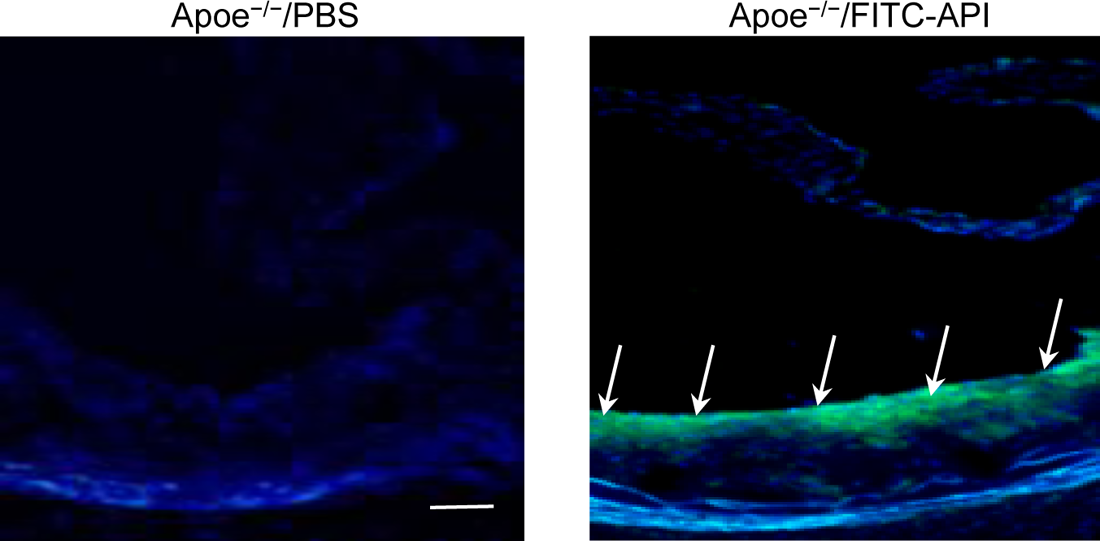
FITC-conjugated API peptide (50 mg/kg, i.v injection) can be targeted to atheroma region in *Apoe^−/−^* mouse model fed WD for 6 weeks. Green color is the FITC concentrated area (FTIC-API). Scale bar: 500 µm.

**Supplemental Figure 14.**
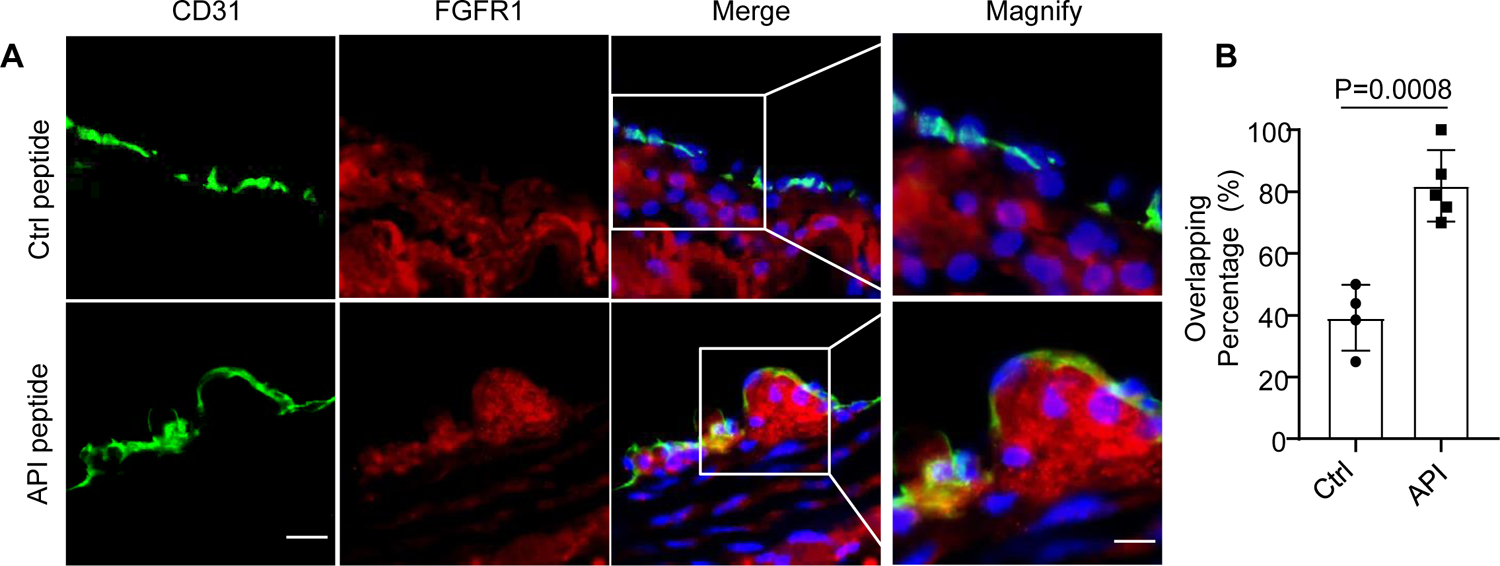
API peptide administration increasing FGFR1 expression in *Apoe^−/−^* mouse model. *Apoe^−/−^* mouse models were administrated with Ctr peptide or API peptide for 12 weeks (50mg/kg, I.V.), BCA was selected for FGFR1 immunofluorescent staining. n=5, P=0.0008, Ctr peptide *vs* API peptide. Scale bar: 20 µm, and magnified: 10 µm.

**Supplemental Table S1:**
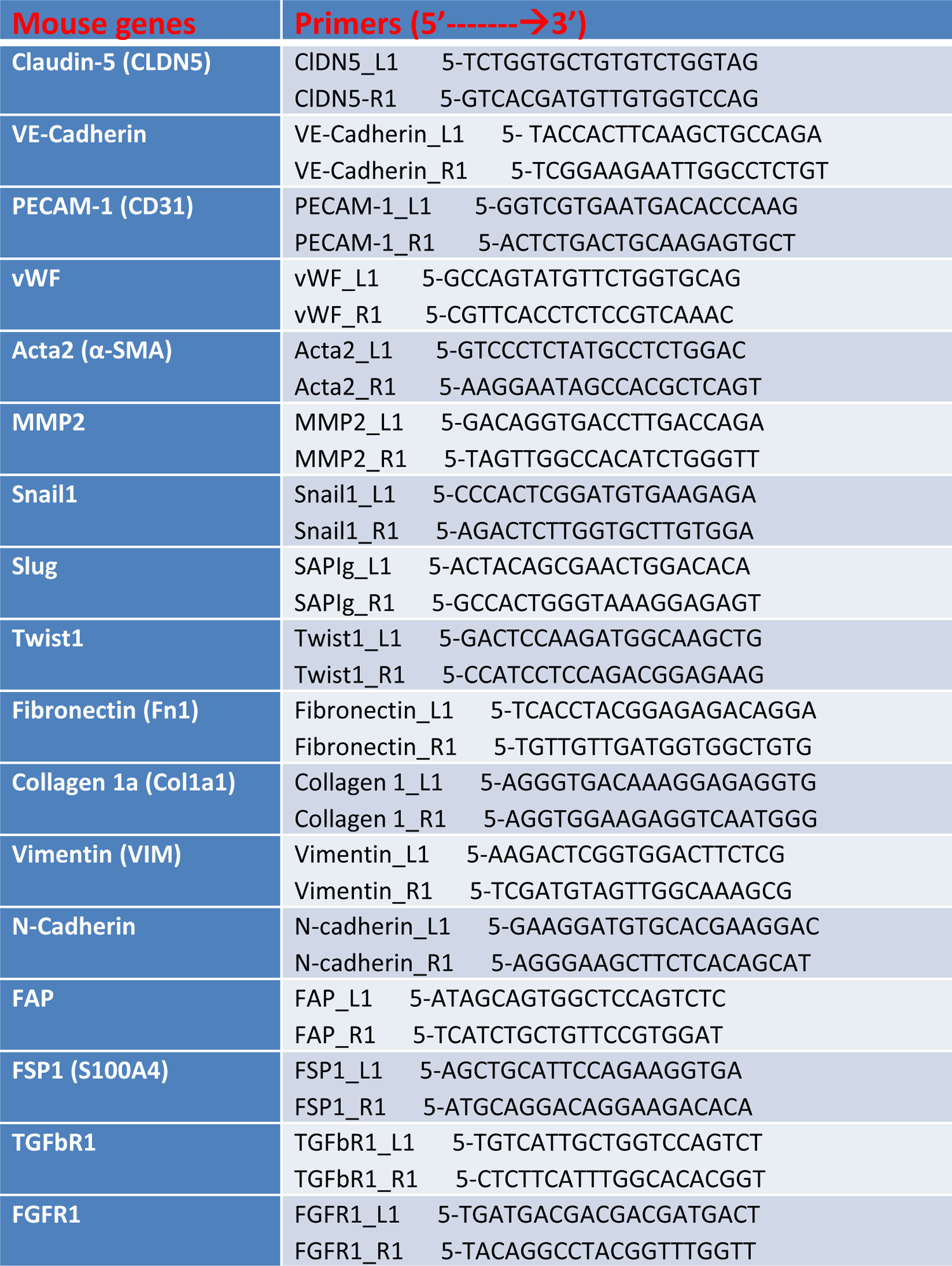
Primers used in the RT-PCR for mouse EndoMT marker genes

**Supplemental Table S2:** Source of antibodies used in this study

| Antibody | Sources | Cat# | Source/<br>Isotype | Applications | Cross<br>activity | MW (kDa) |
| --- | --- | --- | --- | --- | --- | --- |
| FGFR1 | Cell signaling | 9740 | Rabbit IgG | WB, IP, IF, IHC, F | H M R Mk | 92 , 120, 145 |
| Smad2 | Cell signaling | 5339 | Rabbit IgG | WB, IP, IF, CHIP, F | H M R Mk | 60 |
| P-Smad2 | Cell signaling | 3108 | Rabbit IgG | WB | H M R Mi | 60 |
| Smad5 | Cell signaling | 12534 | Rabbit IgG | WB, IP, CHIP | H M R Mk | 60 |
| P-Smad1/5 | Cell signaling | 9516 | Rabbit | WB, IF, F | H M R | 60 |
| Phalloidin-594 | Abcam | ab176757 | MS | IF (red) | M H |  |
| Snail | Cell signaling | 9585 | Rabbit IgG | WB, IP, IF, F | H M | 30 |
| N-Cadherin | Cell signaling | 4061 | Rabbit | WB | H M R Mk | 140 |
| Vimentin | Dako | M7020 | MS | IF | H M |  |
| α-SMA-cy3 | Sigma | C6198-.2ml | MS | IF | H M | |
| VE-Cadherin | Santa Cruz | Sc-9989 | MS | IF | M H |  |
| MYH11 | Abcam | ab224804 | Rabbit | IF, WB | M R H | 42 |
| SM22 (TagIn) | Abcam | ab14106 | Rabbit | IF, WB | M R H | 23 |
| Calponin | Abcam | ab46794 | Rabbit | IF | M R H |  |
| CD31 | BD<br>Pharmingen | 550274 | Rat | IF | M H R |  |
| CD68 | Santa Cruz | Sc-20060 | MS | IF | M H |  |
| Epsin 1 | Home made | n/a | Rabbit | WB, IP, IF | H M | 75~80 |
| Epsin 2 | Home made | n/a | Rabbit | WB, IP, IF | H M | 65~70 |
| HA.11 | BioLegend | 901502 | MS | WB, IP | all | n/a |
| Flag | Cell signaling | 14793 | Rabbit IgG | WB,IP,IHC,IF,F,ChIP | all | n/a |
| α-Tubulin-c | DSHB | 12G10 | MS | WB | M R, etc | 50 |
| β-actin | Cell signaling | 4967 | Rabbit | WB | H M R, etc | 45 |
| GAPDH | Santa Cruz | Sc-166545 | MS | WB | H M | 37-43 |

Supplemental Table S2: Source of antibodies used in this study
| Alexa Fluor 488 (Green) | Alexa Fluor 594 (Red) | Alexa Fluor 647 (Blue) |
| --- | --- | --- |
| Donkey anti-Mouse (A21202) | Donkey anti-Mouse (A23744) | Donkey anti-Mouse (A31571) |
| Donkey anti-Rabbit (A21206) | Donkey anti-Rabbit (A21207) | Donkey anti-Rabbit (A31573) |
| Donkey anti-Rat (A21208) | Donkey anti-Rat (A21209) | Goat anti-Rat (A21247) |
| Donkey anti-Goat (A11055) | Donkey anti-Goat (A11058) |  |

## Notes

### Competing Interest Statement

The authors have declared no competing interest.

### Summary of Updates

Correct author name

